# A metastable contact and structural disorder in the estrogen receptor transactivation domain

**DOI:** 10.1101/332205

**Authors:** Yi Peng, Shufen Cao, Janna Kiselar, Xiangzhu Xiao, Zhanwen Du, An Hsien, Soobin Ko, Yinghua Chen, Prashansa Agrawal, Wenwei Zheng, Wuxian Shi, Wei Jiang, Lin Yang, Mark R. Chance, Witold K. Surewicz, Matthias Buck, Sichun Yang

**Author notes:** Co-first author (Yi Peng, Shufen Cao).

## Abstract

The N-terminal transactivation domain (NTD) of estrogen receptor alpha, a well-known member of the family of intrinsically disordered proteins (IDPs), mediates the receptor’s transactivation function to regulate gene expression. However, an accurate molecular dissection of NTD’s structure-function relationships remains elusive. Here, using small-angle X-ray scattering (SAXS), nuclear magnetic resonance (NMR), circular dichroism, and hydrogen exchange mass spectrometry, we show that NTD adopts a mostly disordered, unexpectedly compact conformation that undergoes structural expansion upon chemical denaturation. By combining SAXS, hydroxyl radical protein footprinting and computational modeling, we derive the ensemble-structures of the NTD and determine its ensemble-contact map that reveals metastable regional and long-range contacts, including interactions between residues I33 and S118. We show that mutation at S118, a known phosphorylation site, promotes conformational changes and increases coactivator binding. We further demonstrate via fluorine-19 (^19^F) NMR that mutations near residue I33 alter ^19^F chemical shifts at residue S118, confirming the proposed I33-S118 contact in the ensemble of structural disorder. These findings extend our understanding of IDPs’ structure-function relationship, and how specific metastable contacts mediate critical functions of disordered proteins.

**Highlights:**

- A compact disorder is observed for the N-terminal domain (NTD) of estrogen receptor
- Multi-technique modeling elucidates the NTD ensemble structures
- Ensemble-based contact map reveals metastable contacts between I33 and S118
- ^19^F-NMR data validate the proposed I33-S118 contact in the IDP

## Introduction

The N-terminal transactivation domain (NTD) of estrogen receptor alpha is critical for the receptor’s constitutive activation via its engagement with coactivator proteins to regulate transcriptional initiation (Lavery and McEwan, 2005; Metivier et al., 2000; Rajbhandari et al., 2012; Sadovsky et al., 1995; Tora et al., 1989; Warnmark et al., 2001). The NTD is an intrinsically disordered protein (IDP), a widely observed structural feature for transactivation domains in various members of the nuclear receptor superfamily (Kumar et al., 2013; Kumar and Thompson, 2012; Li et al., 2012) as well as many other transcription factors such as p53 (Dawson et al., 2003; Lee et al., 2010). Due to the functional significance of the NTDs, they are of interest for drug targeting. In particular, drug candidates that bind an intrinsically disordered region in the NTD of the androgen receptor (AR) have been developed (De Mol et al., 2016; De Mol et al., 2018; Myung et al., 2013). However, understanding a drug’s mechanism of action or the rational of targeting the NTD usually requires in-depth insights into their molecular features. Such knowledge has been hindered by the difficulties in characterizing the structural ensembles of NTDs in particular and IDPs in general. In this paper, we specifically attempt to understand the structure-function relationships of the NTD of the estrogen receptor alpha (ERα) with regard to the structural features that determine its transactivation function.

Current knowledge about the ERα-NTD is limited due to the technical challenges of characterizing IDPs in general (Oldfield and Dunker, 2014; van der Lee et al., 2014). Only limited structural information is available for ERα-NTD from circular dichroism (CD) spectra and chemical shifts from nuclear magnetic resonance (NMR) spectroscopy (Rajbhandari et al., 2012; Warnmark et al., 2001). These have demonstrated that that the isolated NTD is largely unstructured with poorly resolved chemical shifts, and with a high content of random coils and minimal helicity. Recently, biophysical techniques such as small-angle X-ray scattering (SAXS), fluorescence resonance energy transfer (FRET), and hydrogen exchange mass spectrometry (HX-MS) have revealed interesting structural features of IDPs (Berlow et al., 2017; Choy and Forman-Kay, 2001; Lindorff-Larsen et al., 2004; Russel et al., 2012), in particular RXRα-NTD (Belorusova et al., 2016), AR-NTD (Tien and Sadar, 2018), and PR-NTD (Kumar et al., 2013). The application of such approaches to ERα-NTD, however, has not been previously reported to date.

To investigate ERα-NTD structural features under native and denaturing conditions, we conducted multiple, in-solution biophysical studies using a recombinant ERα-NTD protein. We first acquired data from SAXS and HX-MS measurements for the native NTD. HX-MS confirms its overall structural disorder, consistent with the observation by CD and NMR, while SAXS data demonstrate that the NTD is more compact than expected. To examine the influence of chemical denaturants on the NTD, we acquired additional SAXS, CD, and ^1^H-^15^N NMR data for chemically-denatured states and showed that the NTD undergoes structural expansion upon chemical denaturation, indicating a denaturant-dependent loss of structural features that drive the native compactness. To gain insights into the structural basis of this compact disorder, we performed all-atom explicit-solvent simulations to create a structural library of NTD conformations, and supplemented macroscopic SAXS data with hydroxyl radical footprinting coupled with mass spectrometry, which reports solvent accessibility for residue side chains (Huang et al., 2015; Takamoto and Chance, 2006; Wang and Chance, 2017). The combination of these complementary techniques was used to refine and assess ERα-NTD ensemble-structure models. We identified long-range interactions involving residues S118 and I33 driving compactness within the ensemble and showed that S118 mutation mediates conformational changes and alters coactivator binding. Furthermore, through the means of fluorine-19 (^19^F) NMR (Chrisman et al., 2018; Didenko et al., 2013; Kitevski-LeBlanc and Prosser, 2012; Li et al., 2010), we show that mutations near residue I33 alter ^19^F chemical shifts at the position of residue S118, thereby confirming the proposed metastable I33-S118 contact in the ensemble. Overall, our multi-technique biophysical approach allowed us to elucidate specific structural features of the ERα-NTD that mediate its transactivation function, and our findings raise important questions as to the potential for metastable contacts to mediate functions for other nuclear receptors and/or IDPs in general.

## Results

### Compact disorder of ERα-NTD

We purified the ERα-NTD segment (SNA-M1-Y184; Fig. 1A and Fig. 1B) via a refolding process from inclusion bodies of bacterial expression and conducted multiple biophysical measurements to confirm the structural disorder and structural features of this construct under native conditions. First, analysis of circular dichroism (CD) and 2D-NMR spectra reaffirms the disordered nature of the native state as observed by Warnwark and colleagues (Warnmark et al., 2001). Negative ellipticity at 200 nm indicates that the NTD has a high content of random coils with minimal helicity (Fig. 1C). Moreover, the ^1^H-^15^N TROSY spectra exhibit a characteristic spectral crowding in a random-coil region around the 8.0 ppm (Fig. 1D) due to the NTD’s limited chemical shift dispersion (Sahu et al., 2014). We confirm the overall lack of ordered secondary structure by HX-MS measurements (Bai et al., 1993; Buck et al., 1994; Craig et al., 2011; Englander and Kallenbach, 1983; Smirnovas et al., 2011). The HX-MS experiments show nearly 100% deuterium incorporation for the native NTD already after 1 min of exchange (Fig. S1). The only region showing less than complete exchange maps to N- and C-terminal peptides 3-12 and 166-175, respectively. Even though this local protection is very small and potentially within the margin of a systematic or experimental error of the HX-MS measurements, it could also be explained by the presence of a small fraction of residual secondary structure under native conditions as indicated by CD data (see below). Overall, these HX-MS data across the protein sequence suggest that this NTD construct is highly dynamic, consistent with a high level of structural disorder and very little ordered secondary structure as observed by 2D NMR and CD spectroscopy.

**Figure 1.**
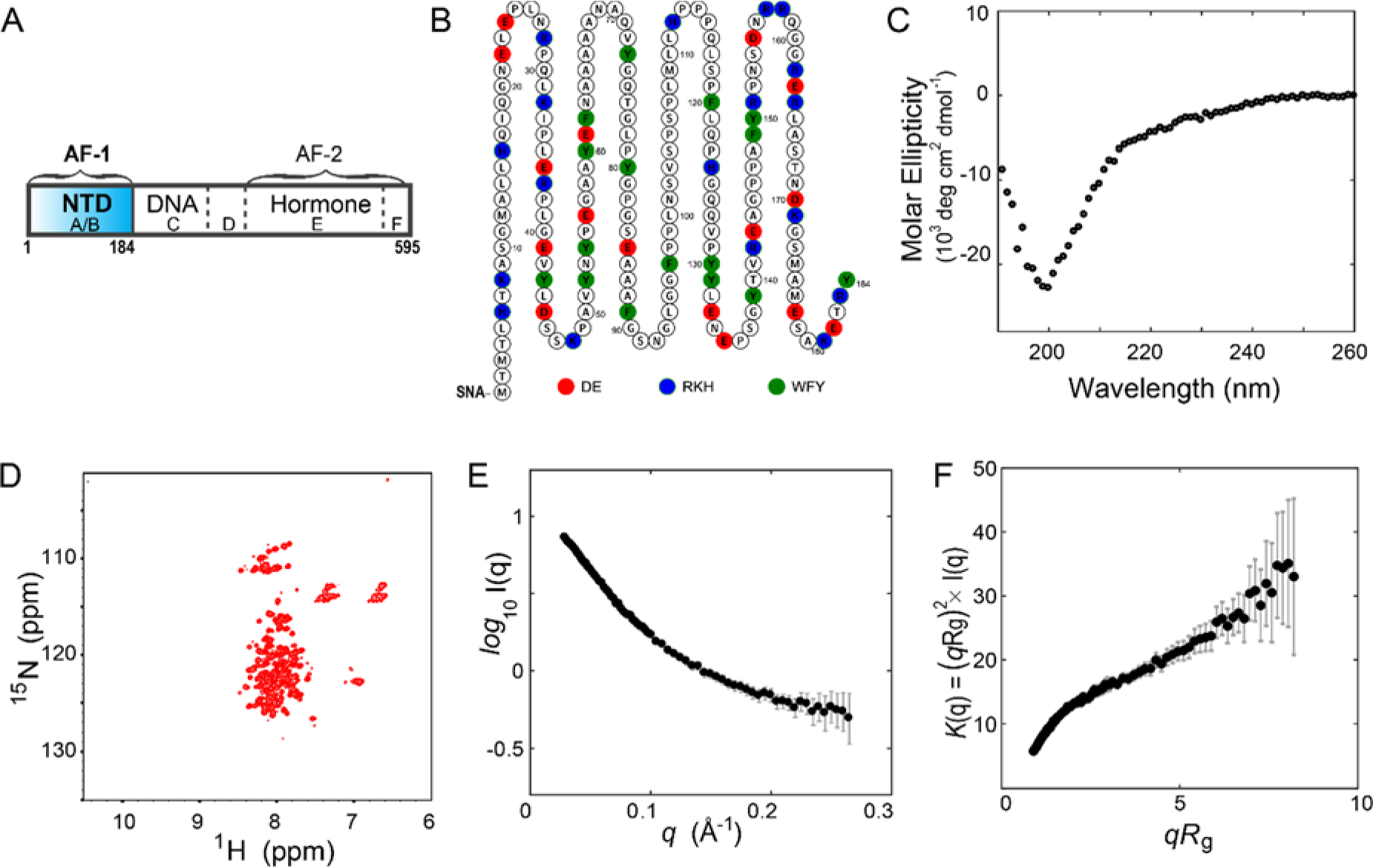
Structural disorder of ERα-NTD. (A) Sub-domains of the ERα. Five domains are the NTD (A/B), a DNA-binding domain (C), a hinge region (D), a ligand-binding domain (E), and a short C-terminal domain (F). (B) The 187-residue sequence. Charged and aromatic residues are highlighted in color. (C) CD spectra for the NTD. (D) Two-dimensional ^1^H-^15^N TROSY spectrum.(E,F) Measured SAXS data yield the R_*g*_ value of 31.0 ± 0.2 Å and its dimensionless Kratky plot shows a structurally disordered characteristic.

We next acquired SAXS data for the NTD in aqueous solution. This SAXS dataset together with related experimental details has been deposited into the Small Angle Scattering Biological Databank (Valentini et al., 2015), with an SASBDB access code SASDEE2 (see also Table S1). The scattering intensity *I*(q) (Fig. 1E) yields a radius of gyration value of *R*_g_^expt^ = 31.0 ± 0.2 Å, using the Sosnick approach (Riback et al., 2017), which is in good consistency with the *R*_g_ values determined using other fitting methods (Table S2). A Kratky plot of *K*(*q*) = (*qR*_g_)^2^\**I*(*q*) shows the characteristics of disordered or unfolded proteins (Fig. 1F), compared to the bell-shaped *K*(q) distribution typically observed in globular well-folded proteins. It is worth noting that many other IDPs approximately followed a power law of *R*_g_^pred^ = 2.54\**N*^0.522^ (Bernado and Svergun, 2012); using this power law, the predicted value is *R*_g_^pred^ = 39.0 Å for the 187-residue NTD, considerably greater than the measured value, even when we consider experimental uncertainties. The apparent difference between *R*_g_^pred^ and *R*_g_^expt^ indicates that ERα-NTD adopts a more compact conformation in its natively disordered state than would be generally expected for IDPs.

To further characterize the protein’s biophysical properties, we calculated the mean hydrophobicity of the NTD at <*H*> = 0.446 (Fig. S2) using the normalized Kyte-Doolittle scale (Kyte and Doolittle, 1982; Wilkins et al., 1999), and determine its mean net charge <*q*> = 0.003 per residue based on the sequence (Fig. 1B). Figure S3 shows that ERα-NTD deviates from the expected combination of <*H*> and <*q*> for IDPs in the two-dimensional H/q space, as it is seen to be to the “right” of Uversky’s proposed boundary line that separates most IDPs and globular folds (Mao et al., 2010; Uversky, 2002). Of note, similar biophysical characteristics are observed for the 130-residue disordered RXRα-NTD which has values of <*H*> = 0.466 and <*q*> = 0.006 (Belorusova et al., 2016), with placement at a similar location in the two-dimensional H/q space. Moreover, we find a balanced charge distribution for ERα-NTD at *f*_+_ = 0.080 and *f*_−_ = 0.086, the fraction of negatively charged (Asp+Glu) and positively charged (Arg+Lys) residues, respectively, consistent with the “weak polyampholyte” regime of IDPs (Das and Pappu, 2013). Overall, these biophysical characteristics of ERα-NTD, at the boundary of native globular and disordered proteins, are consistent with the compact-disordered conformation that the NTD adopts.

### Structural expansion by chemical denaturation

To examine the effect of chemical denaturants on the NTD, we acquired CD data for the NTD in the presence of guanidine hydrochloride (GdnHCl), a chemical denaturant that can disrupt interactions driving the compactness. The minimal ellipticity at 222 nm observed for the native protein is reduced in the presence of GdnHCl (Fig. 2A), indicating the existence of residual secondary structure in the native state. The ^1^H-^15^N TROSY NMR spectra also show increases in structural disorder upon addition of chemical denaturants (Fig. 2B). In the latter experiment, GdnHCl was replaced by urea, a denaturing agent that is more compatible with NMR experiments. By overlapping the TROSY spectra for the NTD in the urea buffer, we observe that NMR peaks are gradually shifted to the downfield region and that the proton’s chemical shift dispersion is narrowed (Fig. 2B top). These NMR peak changes in the presence of a denaturant, together with reduced CD signal at 222 nm, indicate an increase in the level of structural disorder as well as a disappearance of residual secondary structure. We next acquired SAXS data for the NTD in the presence of 3 M and 6 M GdnHCl. Compared with the native *R*_g_^expt^ value of 31.0 Å, the protein exhibits *R*_g_^expt^ values of 38.0 Å and 43.4 Å in the presence of 3 M and 6 M GdnHCl, respectively (Fig. 2C). The latter *R*_g_^expt^ value for the denatured NTD is consistent with those reported for chemically-denatured proteins via a power law (Kohn et al., 2004), by which we estimate a *R*_g_^pred^ value of 44.0 Å for the denatured NTD. This denaturant-dependent size change for the NTD is also consistent with other IDP polypeptide chains that expand with increasing denaturant concentration (Borgia et al., 2016). In this case, however, the native NTD protein is more compact than expected for a typical IDP, and the structural changes seen in NMR and CD upon denaturation apparently disrupt interactions driving the compact state.

**Figure 2.**
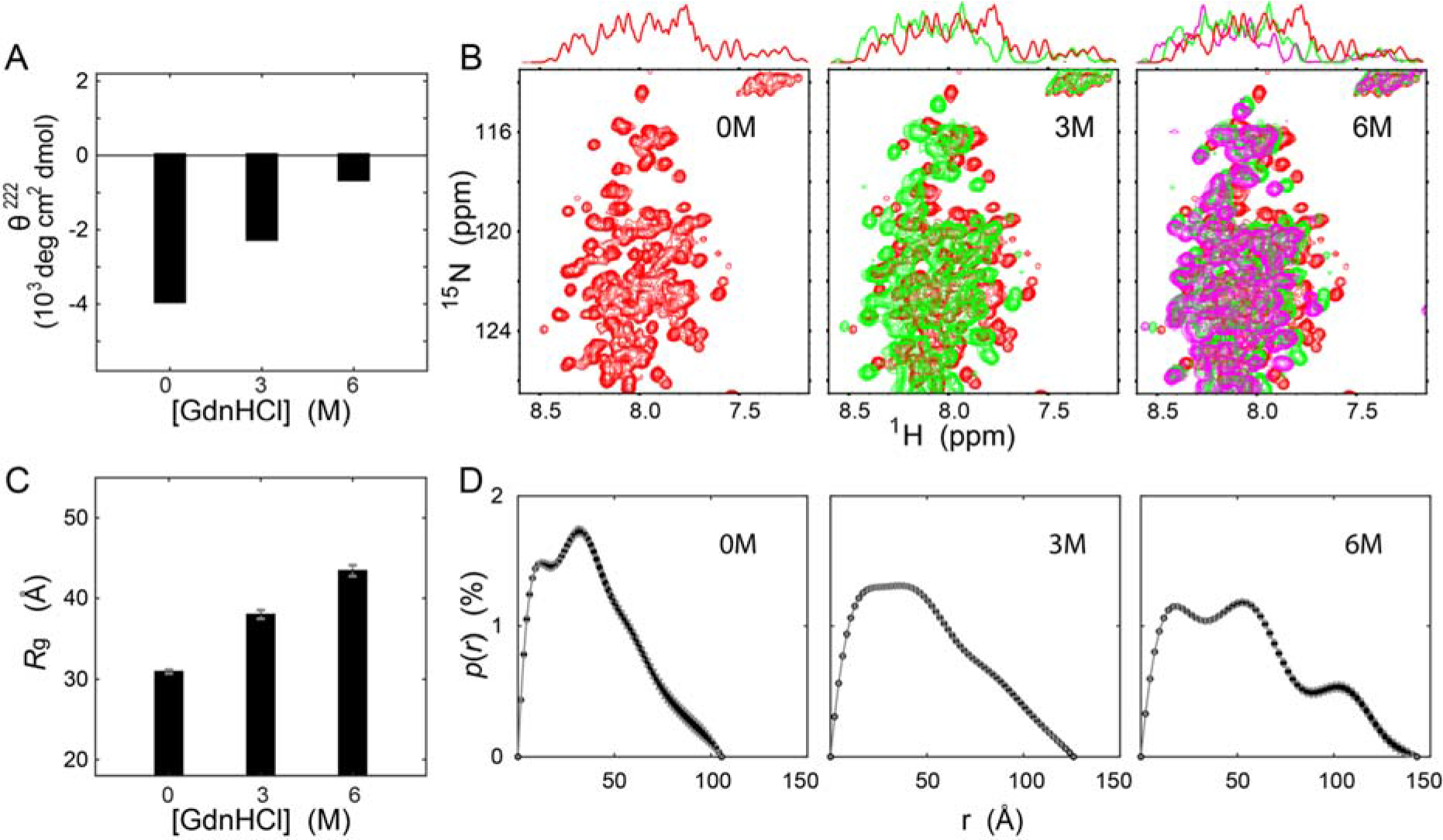
Structural expansion by chemical denaturation. (A) CD signal at 222 nm for ERα-NTD in the presence of 0 M, 3 M, and 6 M GdnHCl, respectively. (B) Two-dimensional ^1^H-^15^N TROSY spectra for the NTD in 0 M (red), 3 M (green), and 6 M (magenta) urea, respectively. On the top is the one-dimensional projection. (C) SAXS-derived *R*_*g*_ values of 31.0 ± 0.2 Å, 38.0 ± 0.5 Å, and 43.4 ± 0.7 Å, respectively, using the Sosnick method (Riback et al., 2017). (D) The pair-wise distance distribution p(r) derived from experimental SAXS data. Calculated using AutoGnom (Svergun, 1992).

To further characterize this variation of the NTD conformation, we used SAXS data to estimate the pair-wise distance distribution p(*r*), a histogram of all pair-wise distances (*r*) between protein atoms. By using an indirect Fourier transformation (Svergun, 1992), we calculated the *p*(*r*) function. For native NTD, this function shows an asymmetric peak below *r* = 50 Å (Fig. 2D), whereas in the presence of 3 M GdnHCl, it is considerably broadened (with a flat plateau), and more so in the presence of 6 M GdnHCl, in which it shows a broad distribution with a long tail up to *r* = 150 Å. The *p*(*r*) analysis shows that the NTD undergoes a considerable expansion from its native compactness with a large-scale stretching out of its polypeptide chain as a result of chemical denaturation.

### Ensemble structures constructed by combining SAXS and FP-PF data

As our studies reveal distinctive features for the native NTD, we attempt to determine the structural basis for the unexpectedly compact state. Given the low-resolution nature of SAXS, however, it is recognized that SAXS data alone are not sufficient to provide details of ensemble-structures. To gain additional information, sequence- or position-specific data from other techniques such as NMR and FRET have been used in conjunction with SAXS for characterizing IDPs (Borgia et al., 2016; Fuertes et al., 2017; Ganguly and Chen, 2009; Krzeminski et al., 2013; Zheng et al., 2016). Here, we supplement SAXS data by hydroxyl radical protein footprinting (FP) coupled with mass spectrometry, where transforming the data to protection factors (PFs) can report solvent accessibility of residue side chains. The value of this approach in multi-technique modeling protein structures and protein-protein interactions has been demonstrated in our previous proof-of-principle studies (Huang et al., 2016) and more recently in the structure determination of a multidomain estrogen receptor complex (Huang et al., 2018). To measure FP-PF data, purified NTD samples were exposed to a focused synchrotron X-ray white beam, where radiolysis-generated hydroxyl radicals react with solvent accessible side chains, and the sites and rates of oxidation are monitored by tandem mass spectrometry MS (Takamoto and Chance, 2006). The extent of modification is quantified as a function of X-ray exposure time (see examples in Fig. S4), yielding a measured rate of footprinting (*k*_fp_) for each site. Dividing the measured rates by those expected for a full accessible site, the intrinsic reactivity with hydroxyl radical (*k*_R_), provides a residue-specific protection factor (PF = *k*_R_/*k*_fp_) (Huang et al., 2015). The natural log of the PF values provides a quantitative residue-specific map on solvent accessibility; high logPF values reflect more protection from solvent (e.g., residues I33 and H124 with logPF = 2.69 and 3.03, respectively), while lower values (e.g., F97 with logPF = 0.68) reflect greater solvent accessibility. Table S3 shows the log PF values for the set of 16 residues that were quantitatively analyzed after Lys-C/Asp-N and/or pepsin digestions. These footprinting protection factor (FP-FP) data provide information about solvent accessibility at the various sites that can be integrated with SAXS data to facilitate ensemble-structure construction for the natively compact-disordered NTD.

We utilized a maximum-likelihood method for ensemble-structure construction that makes use of a structural library of NTD conformations in conjunction with SAXS and FP-PF data, following the general approaches available in the literature (Bernado et al., 2005; Borgia et al., 2016; Brookes and Head-Gordon, 2016; Roux and Weare, 2013; Rozycki et al., 2011; Schwalbe et al., 2014). To generate the conformational library, various computational methods ranging from coarse-grained to simplified all-atom models have been developed (Ghavami et al., 2013; Krzeminski et al., 2013; Ozenne et al., 2012; Vitalis and Pappu, 2009; Zhang and Chen, 2014). Due to the prior knowledge about the NTD compactness, here we used an accumulative set of 35-μs all-atom explicit-solvent molecular dynamics (MD) simulation trajectories (Methods). The resulting pool of NTD conformations is combined with and compared against experimental SAXS and FP-PF data simultaneously to construct the best-fit ensemble-structures. The goodness of fit between the ensemble-calculated and experiment data is assessed by two unit-less functions χ^2^ (Eq. 1) and φ^2^ (Eq. 2), each measuring the difference between calculated and experimental SAXS and FP-PF data, respectively. To avoid potential over-fitting, we applied a maximum-likelihood method (Eq. 3, Eq. 4, and Fig. S5). As such, a broad probability distribution {*P*_*i*_} is determined for the ensemble-structures. It is clear that while the MD simulations were able to sample a broad range of conformations (e.g., with a wide *R*_g_ distribution; Fig. S6A), the ensemble-fitting against experimental SAXS and FP-PF data only selects a subset of sampled conformations that better match experimental SAXS and footprinting-related SA values (Fig. S6B). Overall, this multi-technique modeling for the NTD resulted in an ensemble of top 10 clusters of structures (Fig. 3A) that contributes to a total of 16% of the entire population and an ensemble of top 100 structures that accounts for about 80% of the population (Fig. 3B).

**Figure 3.**
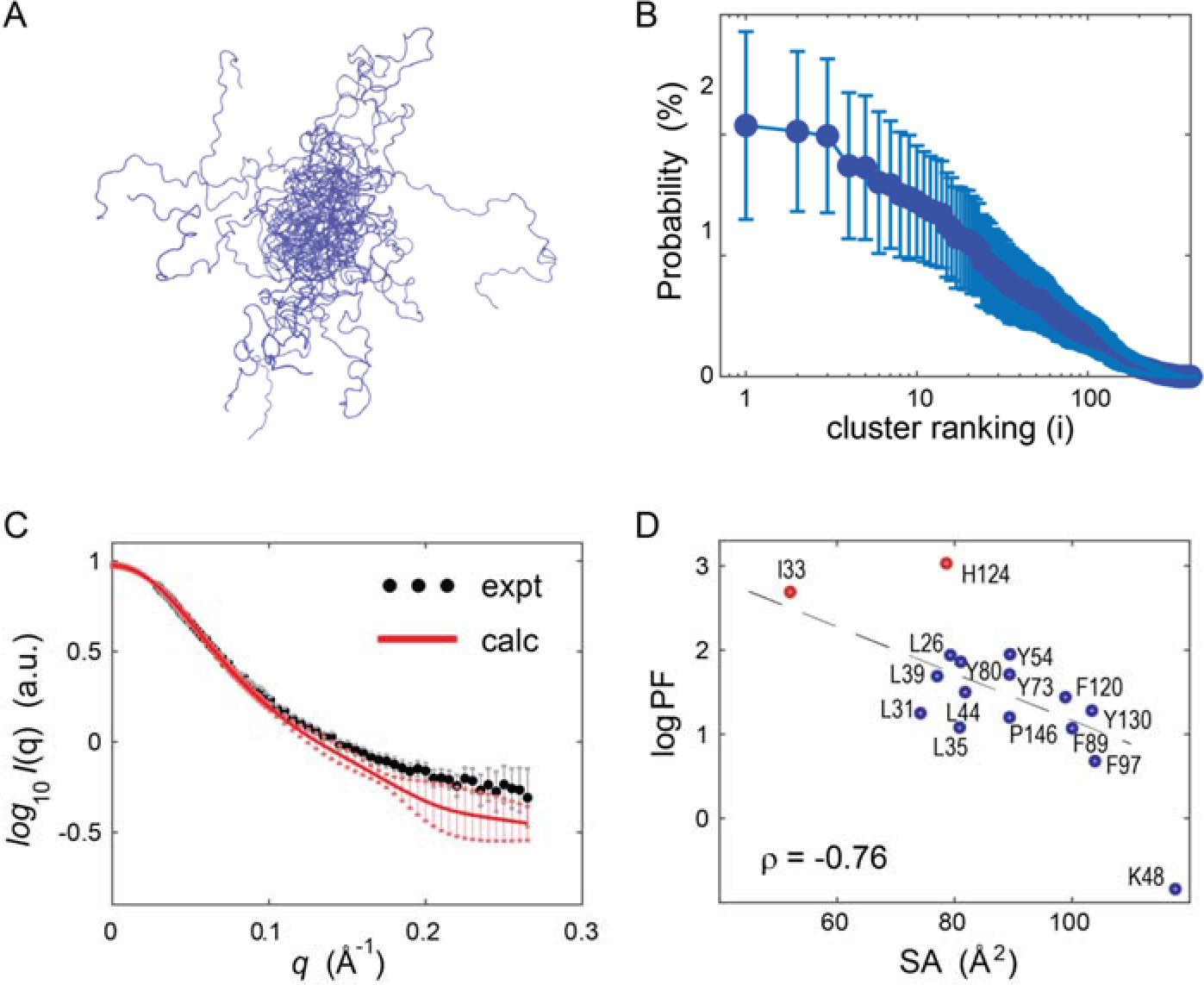
Ensemble-structure construction for the native compact-disordered NTD. (A) Ensemble-structures constructed by integrating SAXS and FP-PF data. Shown is the ensemble of the top 10 structures. (B) Probability distribution for the ensemble-structures. The error bars were the standard deviations from nine sets of probability distributions calculated using a same set of conformations. (C) Measured and calculated SAXS profiles. The calculated SAXS profile (in red) is the average of ensemble-structures. The SAXS data were deposited into the SASBDB databank (https://www.sasbdb.org/data/SASDEE2). (D) Measured logPF values and ensemble-based solvent-accessible surface areas (SAs). A set of 16 residue side-chains were probed and shown.

Comparison of the ensemble-calculated and experimental SAXS profiles shows a reasonably good agreement (χ^2^ = 1.7 as defined in Eq. 1; see Fig. 3C). In addition, a weighted average for side-chain solvent-accessible surface area (SA) is estimated for each residue and plotted against its corresponding logPF value (Fig. 3D). The SA vs. logPF plot has a correlation coefficient of −0.76 (*p*-value = 6.3×10^−4^ for the set of 16 residues), suggesting reasonable agreement between the derived ensemble-structures and experimental footprinting data. Previous correlation analyses of this kind have been performed on globular folded proteins, which typically exhibit a wide range of logPF values. However, most of the observations here have logPF values from 1 to 2 (SA values of 80-110 Å^2^), and have few values at the extremes of being either highly solvent-protected or highly solvent-accessible (e.g., upper left or lower right quadrants). In order to remove any potential bias due to residue size, we normalized the SA data into the fractional SA (fSA) values, using the maximum possible SA for the residue (Tien et al., 2013). In a re-plotted fSA vs. logPF in Fig. S7, 12 out of the 16 points (circled) are clustered in the “middle” of the plot in a restricted range. This contrasts with globular folded proteins that exhibit as many residues in the “middle” range as in the high/low fSA regions (Gustavsson et al., 2017; Kaur et al., 2015). This is presumably because the existence of a balanced distribution, including both high (e.g., fSA > 0.8) and low (e.g., fSA < 0.4) solvent-accessible residues, correlates with structural stability and maintains the overall fold for globular folded proteins. By comparison, the majority of NTD residues are in an intermediate range of fSA (and logPF) values, consistent with its dynamic but compact nature. This relatively narrow fSA distribution is also consistent with the modest chemical shift dispersion observed in NMR spectra and may reflect a structural signature of this NTD. Retrospectively, this broad distribution for the ensemble is also consistent with the typical behavior of disordered proteins, where an ensemble of structures is often required to characterize the structural properties of IDPs.

We next performed an ensemble contact-map analysis for the NTD ensemble-structures (Methods). The resulting contact-map shows that the NTD has multiple medium and long-range contacts formed (within residues 11-75), pointing to a “collapsed” segment (for a portion of the distribution) at its N-terminus, compared to its C-terminus (Fig. 4A). This regional collapse is consistent with the higher hydrophobicity of its N-terminal sequence than the C-terminal (Fig. S2). Moreover, long-range interactions are formed (e.g., frequently observed in the ensemble) between two 5-residue peptides, each centered at I33 and S118 (marked by an arrow in Fig. 4A and illustrated in Fig. 4B). Residue I33 is predicted to have a SA value of 52 Å^2^ from the ensemble-structures (Table S3), indicating that I33 is more protected from solvent exposure than the typical or average member of the ensemble. To gain insight into the nature of these conformations and their distributions, we examined the explicit structural distribution of I33, compared to L44, which has a similar/compatible side-chain size but with a higher SA value indicating much greater solvent exposure. It can be seen that based on the top 100 structures (Fig. 4C), I33 has far more ensemble members with SA values below 50 Å^2^, compared to L44 that has a greater number of conformations above 100 Å^2^. I33 is on average solvent-protected to a greater degree, presumably due to its “metastable” interaction with S118 (see below) that tilts the conformational distribution towards greater protection, compared to L44. While the interaction itself is presumably relatively weak compared to tertiary contacts for a globular well-folded protein, the metastable nature of this long-range interaction could provide a critical structural role driving the function of the NTD (Wright and Dyson, 2015).

**Figure 4.**
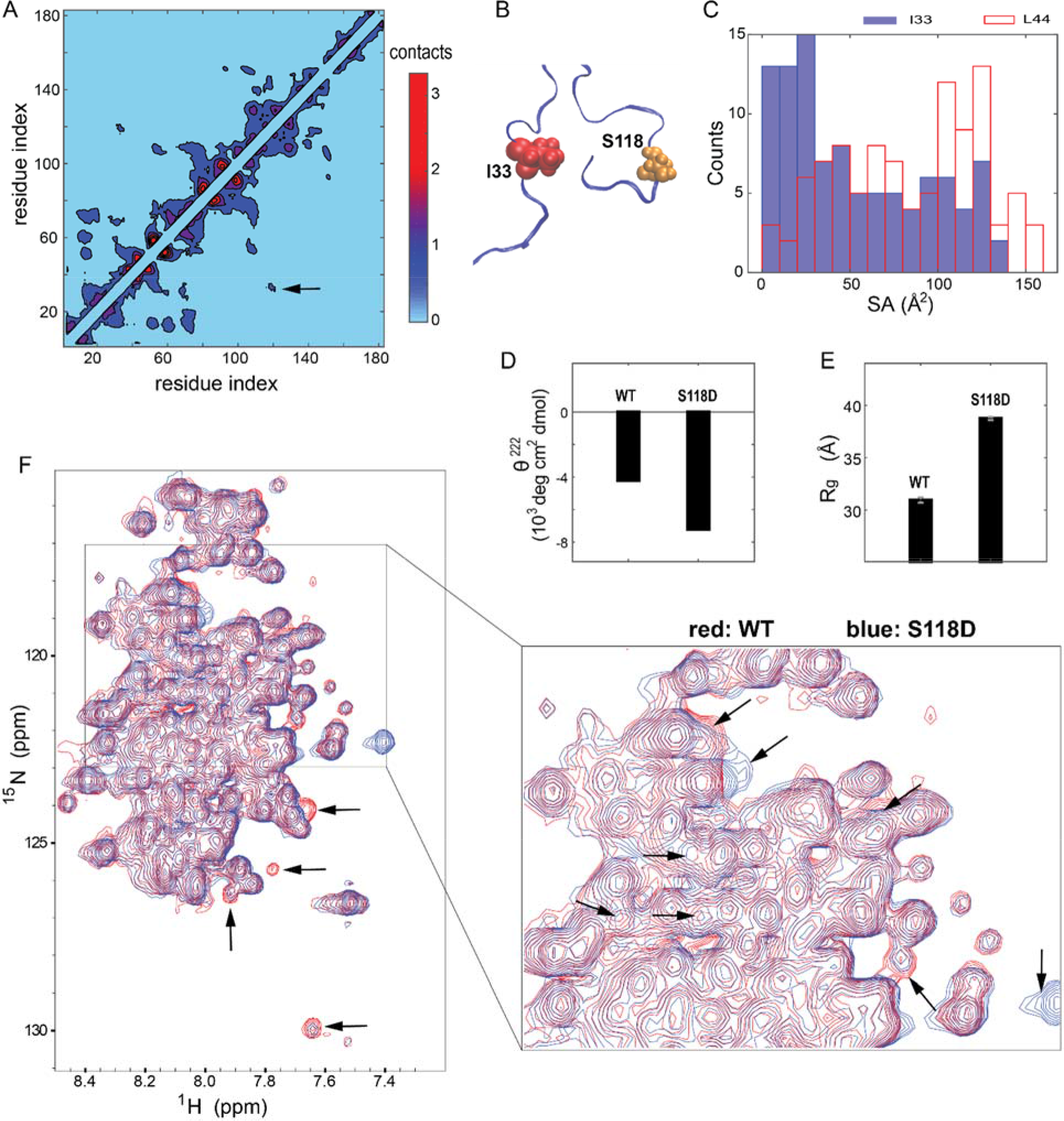
Ensemble-contact map and mutation-induced conformational changes. (A) Contact map shows that its N-terminal region (e.g., residues 11-75) has more contacts formed than its C-terminal. The color-bar indicates the contact number, where each contact is ensemble-weighted (Methods). The metastable I33-S118 contact is marked by an arrow, where three of the top 10 structures from the ensemble make the contact. (B) Cartoon illustration of I33-S118 interaction. (C) Histogram for SA values for residues I33 and L44 from top 100 structures of the ensemble. (D) CD signal at 222 nm for S118D as well as wild-type (WT). (E) The *R*_*g*_ value of 38.7 ± 0.2 Å for mutation S118D, compared to the *R*_*g*_ value of 31.0 ± 0.2 Å for the wild-type (WT). Related SAXS data shown in Fig. S8. (F) Overlap of ^1^H-^15^N TROSY spectra between WT (red) and mutant S118D (blue). Peak changes/shifts marked by arrows.

To examine the functional role of S118 and to test its structural interaction predicted by our ensemble-structures, we mutated the non-charged S118 residue to a charged residue Asp (D) (i.e., S118D), as a phosphomimetic mutation. The ellipticity at 222 nm shows that S118D increases the content of ordered secondary structure (Fig. 4D). To gain insight into the impact on the overall protein, we further acquired SAXS data for the S118D mutant, yielding the *R*_g_^expt^ value of 38.7 Å (Fig. 4E and Fig. S8), which is greater than that of the wild-type (*R*_g_^expt^ = 31.0 Å) and smaller than the denatured state (*R*_g_^expt^ = 43.4 Å for the NTD in the presence of 6 M GdnHCl). Mutation-dependent conformational changes are also confirmed by peak shifts and appearance/disappearance in the ^1^H-^15^N TROSY spectra of S118D (Fig. 4F; arrows). While detailed NMR assignments to identify specific residues involved will be the subject of future studies, these considerable changes involving at least 12 spectrum peaks indicate that S118D mutation leads to modest conformational changes that spread out well beyond the mutation site itself, in accord with the sizeable *R*_g_ change as observed by SAXS. Overall, these data collectively show that the mutation propagates its structural impact and physically extends the polypeptide chain, presumably by destabilizing S118-mediated long-range contacts, while counterintuitively resulting in a gain of local secondary structure.

We also examined the impact of the S118D mutation on the NTD binding to TBP, a general transcription factor TATA box-binding protein, known as an ER coactivator in the transcriptional machinery (Louder et al., 2016; Sadovsky et al., 1995). We estimated a weak-to-modest binding affinity of 53 ± 12 μM between the wild-type NTD and TBP, using surface plasmon resonance (SPR) spectrometry (Fig. S9), consistent with the previously reported binding affinity in the micromolar range (Warnmark et al., 2001). The NTD-TBP binding is further demonstrated by NMR experiments, which show more than 15 peak shifts and appearance/disappearance in the ^1^H-^15^N spectra of the wild-type (WT) upon addition of TBP (Fig. S10A). Compared to the WT, however, the addition of TBP leads to a greater extent of changes in the ^1^H-^15^N spectra of the S118D mutant, with a set of 23 peak changes (Fig. S10B; arrows), indicating that S118D increases coactivator binding as a result of the conformational changes. This increased coactivator binding is further supported by the SPR measurements, where S118D increases the TBP-binding affinity more than ten-fold to *K*_d_ = 3.9 ± 0.4 μM (Fig. S11), compared to the WT’s *K*_d_ = 53 ± 12 μM. Together, these data show that the S118D mutation induces sizable conformational changes within the NTD and further increases the coactivator binding.

### Metastable contact in the IDP validated by ^19^F-NMR

To test the validity of the proposed I33-S118 interaction, we conducted ^19^F-NMR studies on the NTD by introducing a cysteine residue at the location of S118 to allow covalently-linked ^19^F labeling with a trifluoromethyl (–CF_3_) group (depicted in Figure 5A,B). The ^19^F labeling was achieved using 3-Bromo-1,1,1-trifluoroacetone (BTFA), a wildly-used labeling reagent in ^19^F-NMR studies of both disordered and well-folded proteins (Didenko et al., 2013; Kitevski-LeBlanc and Prosser, 2012; Li et al., 2010), with a more recent application to the nuclear receptor PPARγ (Chrisman et al., 2018). Hereafter, we refer to the BTFA-labeled NTD S118C protein variant as NTD^S118C^-BTFA. Because there is no cysteine in the native NTD, ^19^F-NMR spectra of the NTD^S118C^-BTFA yielded a single peak in each ^19^F-NMR spectrum (Figure 5C, left). Of note, the relative exposure to the solvent of the BTFA probe can be qualitatively determined by the ^19^F peak shift as a function of D_2_O concentration, an effect known as D_2_O-sovlent induced isotope shift (Chrisman et al., 2018; Kitevski-LeBlanc and Prosser, 2012). Here, the “wild-type” form of the NTD^S118C^-BTFA was observed with a peak shift Δδ of 7.5×10^−2^ ppm between 10% and 50% D_2_O. We next determined the effect of mutations near residue I33 via two mutations: a single-point mutation of L31A and a triple mutation of K32A/I33A/P34A. The ^19^F-NMR spectra of the L31A showed a negligible difference in the peak shift Δδ, compared to the wide-type NTD^S118C^-BTFA (Figure 5C, middle). In contrast to L31A, as a negative control, the triple mutant K32A/I33A/P34A was observed with a Δδ value of 9.2×10^−2^ ppm (Figure 5C, right), which is a 23% increase from the wild-type. Of note, while the D_2_O itself causes a small chemical shift, the focus here is on the Δδ difference between 10% and 50% D_2_O, as opposed to the absolute chemical shift with a specific D_2_O-concentration buffer. Given the large sequence separation between I33 and S118, the K32A/I33A/P34A mutations are observed with an unexpectedly impact on the surroundings of residue S118. These ^19^F-NMR data of the wide-type and the triple mutant provide a strong evidence that the mutations disrupt the I33-S118 contact, thereby increasing the solvent exposure of the ^19^F probe at residue S118. Given a wide range of conformations available for the IDP and the metastability of the long-range contact, it is difficult for a majority of biophysical techniques to achieve a validity test on such transient contacts. Here, the one-dimensional ^19^F-NMR spectroscopy, due in part to the natural abundance of fluorine-19, offers a highly-sensitive tool to detect subtle changes associated with the NTD contact. As such, our ^19^F-NMR data together with site-directed mutagenesis studies confirm the proposed metastable contact between residues I33 and S118 of the NTD ensemble.

**Figure 5.**
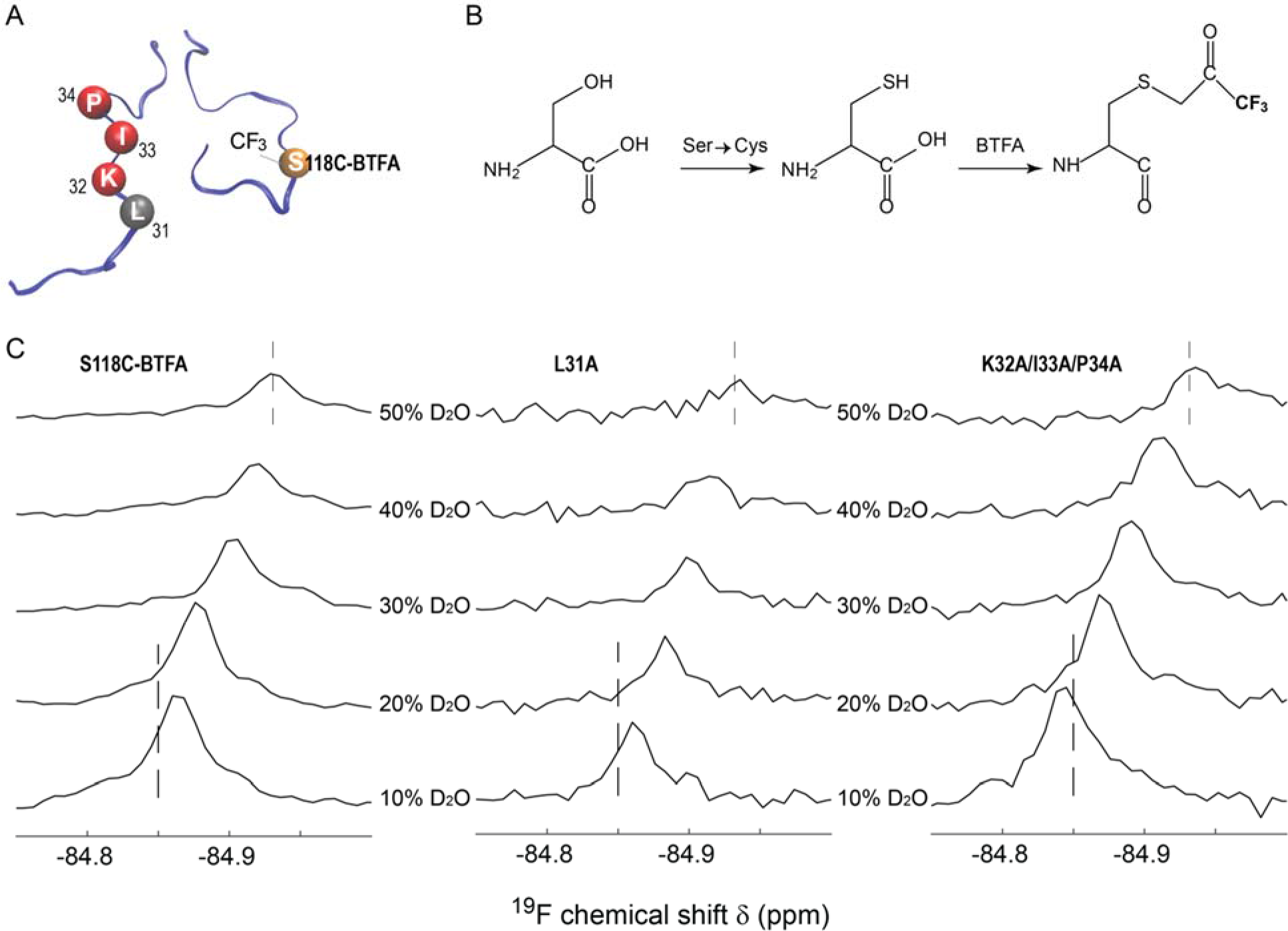
Mutations near residue I33 alter the fluorine-19 chemical shift at residue S118. (A) Location of the BTFA (3-Bromo-1,1,1-trifluoroacetone) at S118. Sites of mutation near I33 (the point mutation L31A and the triple mutation K32A/I33A/P34A) are also labeled. (B) Schematic of the NTD^S118C^-BTFA labeling where fluorine-19 (–CF_3_ group) is covalently linked to the cysteine of mutant S118C in a one-step chemical reaction. (C) ^19^F-NMR spectra of the wild-type, L31A and K32A/I33A/P34A of the NTD^S118C^-BTFA under different D_2_O concentrations. Compared with the wild-type NTD^S118C^-BTFA, ^19^F chemical shifts are observed with a negligible change for mutant L31A, but with a 23% increase for the K32A/I33A/P34A (between 10% and 50% D_2_O), as a result of the effect of D_2_O-solvent induced isotope shift. Dash lines were used for visualization only.

## Discussion

The NTD accounts for a large portion of the total transcriptional activities of most nuclear receptors in terms of transactivation function. While it is recognized early on that ERα-NTD is structurally disordered, little is known about its molecular details, compared to extensive knowledge available for other regions of the receptor such as the DNA-binding domain and the hormone-binding domain (Fig. 1A). This study provides a comprehensive characterization of the NTD’s rather distinctive structural features for both native and chemically-denatured states. Our biophysical measurements show that the NTD is mostly disordered as revealed by CD and NMR, although SAXS data point to considerable compactness, in accord with the modest hydrophobicity and nearly-zero net charge that is more common for globular proteins. The ensemble structures were observed to undergo a loss of residual secondary structure and a structural expansion from a narrow single-peak distance distribution in the native state to a broad distribution in chemically-denatured states, indicative of melting of pre-existing structural features associated with the compactness and/or helicity. Of note, this phenomenon of a large expansion with >40% increase in *R*_g_ from 31 to 43 Å was observed for ERα-NTD only; a generalization to and comparison with other IDPs such as RXRα-NTD and AR-NTD has not yet been attempted.

The ensemble-structures constructed by multi-technique modeling, which combines SAXS and FP-PF data together with a library of simulation-generated conformations, provide a rare opportunity to observe the molecular details of ERα-NTD. We acknowledge that this modeling process cannot provide a unique solution to understanding the ERα-NTD structures in part due to the ensemble-average and/or conformer-coexistence nature of both SAXS and FP-PF data (*i.e.*, footprinting protection factors). We further note that several residues had extreme logPF values not explained by the modeling. In particular, H124 was observed to be the most protected, but our ensemble-structure models could not explain its high degree of solvent protection (Fig. 3D), which points to next-steps for future ensemble refinements. Nonetheless, given the strong evidence that validates our constructed ensemble-structures for ERα-NTD, our multi-technique approach could be applied for the structure determination of other IDP systems such as RXRα-NTD.

In a broader context, IDPs are increasingly appreciated as functionally important for coactivator recruitment of protein-protein interactions involved in cellular signaling and transcriptional regulation. Typical IDPs are viewed as a diverse population of interchangeable structures or conformers that co-exist in a concerted equilibrium. Here, the ensemble-structure studies for ERα-NTD add to our understanding of the range of IDPs’ structure-function capability. A large portion of the ensemble population for ERα-NTD favors a regional collapse at the N-terminus contributing to the overall compactness. Moreover, a subset of the population, roughly three out of the top 10 structures in the ensemble, adopt long-range contacts near I33 and S118 that presumably tilt the dynamic equilibrium in favor of more compact conformers. The extended NTD ensemble, accompanied by the increase of ordered secondary structure, possesses structural elements or flexibility that permits the IDP to interact more productively with the folded TBP protein, resulting in stronger coactivator binding. Functionally, S118 is a known phosphorylation site that modulates the receptor’s transcriptional activity (Chen et al., 2000), and an increase in the level of S118 phosphorylation has been associated with endocrine resistance via direct recruitment of coactivators (Kato et al., 1995). As a phosphomimetic mutation, S118D directly links a phosphorylation-driven shift in the NTD conformational equilibrium in favor of coactivator recruitment, suggesting the role of residue S118 as a “structural inhibitor” and further providing a molecular basis for understanding S118-associated endocrine resistance in breast cancer. S118 phosphorylation is also critical for association with a cis/trans isomerase pin1 as observed in a truncated NTD segment, while mutation of S118 to alanine prevents the association (Rajbhandari et al., 2012). Such mutation-induced modulation is also observed in other IDPs such as the transcription factor HIF-1α’s transactivation domain (Berlow et al., 2017) and a disordered PAGE4 protein (Kulkarni et al., 2017). More strikingly, despite the “metastable” nature of the I33-S118 contact, mutation near residue I33 considerably alters the ^19^F chemical shifts at the position of S118C at a distance along the amino acid sequence, thereby confirming the structural articulation of this long-range contact observed in the IDP ensemble-structures. Because there exist a wide range of conformations for the IDP and it is difficult to directly visualize any ensemble-averaged features of these conformations, our contact map analysis enables an ensemble-structure approach for studying disordered proteins by making testable predictions on the I33-S118 contact being the case in point. In conclusion, as we note above that the hydrophobicity and mean net charge of RXRα-NTD is similar to that of ERα-NTD, our findings suggest that a comprehensive characterization of RXRα-NTD and/or other NTDs by these approaches may reveal additional examples of metastable contacts in the context of IDPs that fine-tune the conformational equilibrium to execute particular molecular functions as a general mechanism of mediating nuclear receptor structure and function.

## Author contributions

S.Y. designed the research. Y.P., A.H., and S.K. expressed and purified the protein. Y.P. performed CD experiments. S.C., Y.P. and M.B. contributed to 2D-NMR experiments. X.X., Y.P., Z.D. and W.K.S. contributed to H/D exchange experiments. S.Y., W.S. and L.Y. collected scattering data. J.K., Y.P., M.R.C. and S.Y. contributed to footprinting experiments. W.J. and S.Y. performed MD simulations. S.Y. performed the integrative modeling. W.Z. contributed to the *R*_g_ analysis. Y.P. and Y.C. performed SPR. Y.P., P.A. and Z.D. contributed to the ^19^F-NMR experiments. All authors edited the manuscript.

## Acknowledgments

We thank Sayan Gupta for assistance with footprinting experiments, Geof Greene for inspiration of this project, Marc Parisien for constructive comments, and Liang-Yuan Chiu and Hao Ruan for assistance with ^19^F-NMR experiments. We also thank anonymous reviewers for constructive suggestions. The work of S.Y. was supported by a NIH grant (GM114056). Use of the synchrotron sources was supported by the U.S. Department of Energy (DE-AC02-98CH10886) and by NIH (EB009998).

## STAR× Methods

### Key Resources Table

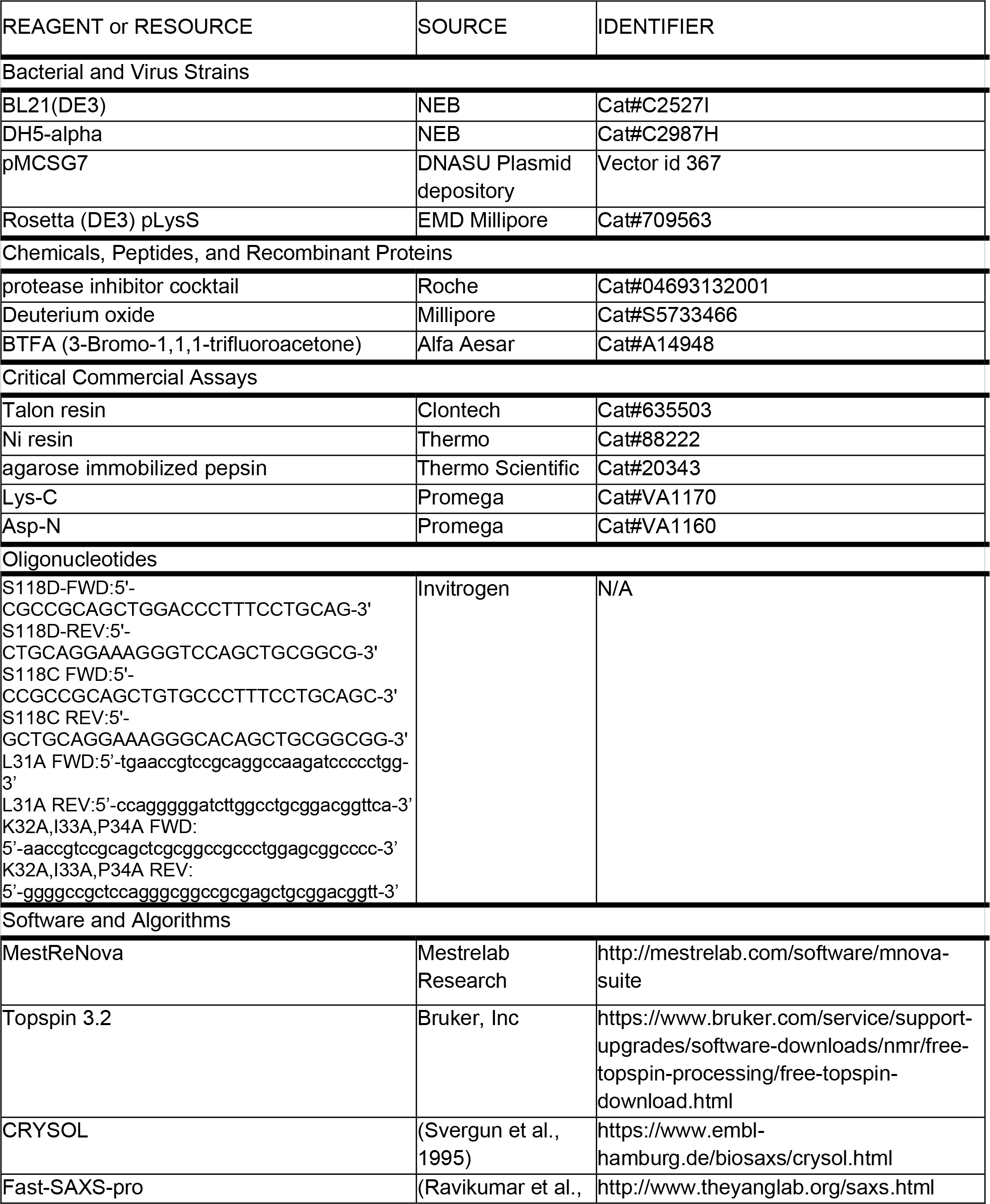
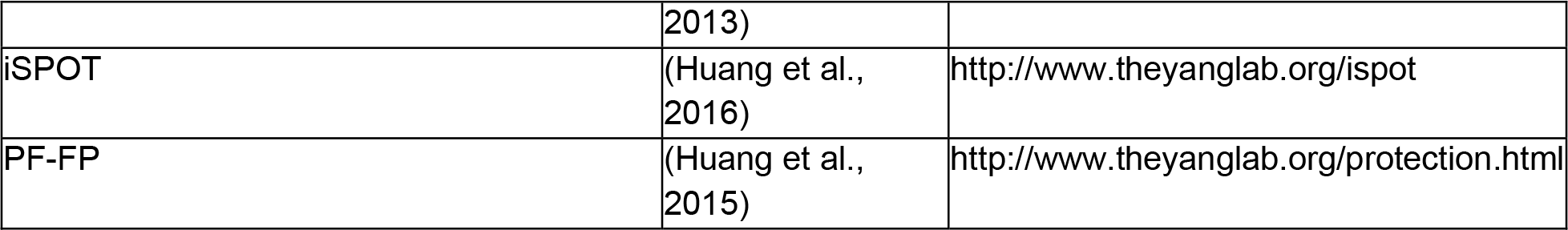

### Contact for Reagent and Resource Sharing

Further information and requests for reagents may be directed to and will be fulfilled by the Lead Contact Sichun Yang.

## METHOD DETAILS

### Plasmid Construct

The sequence of the N-terminal domain (NTD) of human ERα (M1-Y184) was inserted into a bacterial expression vector pMCSG7 containing an N-terminal 6-Histidine tag designed for Ni-NTA affinity purification. Verification of this expression vector was performed for DNA sequencing. His-tag was cleaved by TEV protease to generate the NTD protein with a short three-residue segment of SNA (from the expression vector) at its N-terminus.

### Protein Expression and Purification

The NTD samples were expressed as inclusion bodies in bacterial BL21 (DE3) cells. Expression was carried out overnight at 18 °C with 0.1 mM IPTG in TB medium after the OD_600_ reached ~0.8. Cells were harvested by centrifugation at 5000 rpm, re-suspended in 50 mM Tris, pH 8.0, 500 mM NaCl with a protease inhibitor cocktail (Roche, Indianapolis, IN). The cells were disrupted using a M110Y microfluidizer (Microfluidics, Newton, MA). The pellet fraction containing the NTD inclusion bodies was solubilized in the buffer of 50 mM sodium phosphate, pH 7.4, 300 mM NaCl, 6 M guanidine hydrochloride, and 5 mM imidazole with 1 mM PMSF. Solubilized inclusion bodies were centrifuged at 18,000×g for 45 min at 4 °C. Cleared supernatant was incubated with Talon resin (Clontech) for 45 min at 4 °C, followed by a wash with a buffer of 50 mM sodium phosphate, pH 7.4, 500 mM NaCl, 5 mM imidazole, and 6 M guanidine hydrochloride. Samples were step-eluted with imidazole at a series of concentrations of 50, 250, and 500 mM. Fractions containing the His-tagged NTD protein were combined and dialyzed against a buffer of 20 mM Tris, pH 7.5, 10 mM NaCl and 0.1 mM PMSF. Refolded NTD proteins were collected and incubated with TEV protease at a TEV:NTD ratio of 1:50 overnight at 4 °C. The second step of affinity purification via Talon resin was used to remove the TEV and uncleaved His-tagged proteins. Final purification was performed using ion exchange via a HiTrap Q 5 mL column (GE Healthcare) and by gel filtration via an Enrich SEC650 column (Bio-Rad).

### Circular Dichroism Spectroscopy

NTD proteins in a buffer of 20 mM sodium phosphate (pH 7.4) were used for the far UV CD spectroscopy measurement at 25 °C on an AVIV 215 spectrometer with a 0.1 cm quartz cell. CD spectra were recorded in the wavelength of 190-260 nm. Each CD spectra were reported after buffer subtraction. Final spectra were averaged over three independent scans.

### Two-dimensional NMR Spectroscopy

Two-dimensional NMR spectra were recorded at 8 °C on a Bruker Avance II 800-MHz spectrometer equipped with a TXI cryoprobe. The NTD protein was prepared in the buffer of 10 mM sodium phosphate, pH 7.2, 100 mM NaCl, 0.5 mM EDTA and 0.1mM PMSF, in the presence of 0 M, 3 M, 6 M urea, respectively, with a final protein concentration of 100 μM. Here, urea was used to replace GdnHCl because of its chemical compatibility for NMR spectroscopy. Two-dimensional ^1^H-^15^N TROSY spectra were collected with 1024×248 (F1×F2) points and 64 scans. In addition, one-dimensional proton spectra were recorded, where the ^1^H chemical shifts of Arginine side chains (at 6.92 ppm and 6.97 ppm) were identified and used as a reference to calibrate the ^1^H-^15^N TROSY spectra. Spectra were processed with NMRPipe (Delaglio et al., 1995) and analyzed with Sparky (Goddard TD, Kneller DG, SPARKY 3, University of California, San Francisco).

### Small-angle X-ray Scattering

A flow-cell setup was used for SAXS measurements at the LiX-16-ID beamline at the National Synchrotron Light Source II of the Brookhaven National Laboratory (Upton, NY). A series of scattering images were recorded each with a 2-sec exposure for both protein and buffer samples. Final scattering intensity was reported after buffer subtraction. The X-ray energy was 13 KeV.

### Amide Hydrogen Exchange with Mass Spectrometry

Purified NTD (4.6 ug/ul) was prepared in 20 mM sodium phosphate buffer (pH 7.4). To initiate the deuterium labeling, 1ul aliquots of purified NTD were added to 19 μl of 10 mM phosphate buffer (pH 7.3) in D_2_O. After incubation at room temperature for different time periods (1 min and 5 min), samples were cooled on ice in a pre-calibrated quenching 0.1 M phosphate buffer (pH 2.4) containing 7 M GdnHCl and 0.1 M TCEP. The solution was diluted 10 times with ice-cold 0.05% trifluoroacetic acid and digested for 5 min with agarose immobilized pepsin (Thermo Scientific). Peptic fragments were separated and analyzed by an LTQ Orbitrap XL mass spectrometer (ThermoElectron, San Jose, CA) as described previously (Smirnovas et al., 2011; Smirnovas et al., 2009). As a result, the percent deuterium uptakes were averaged for the triplicate samples of NTD under each condition to obtain the mean percent deuterium incorporation.

### Hydroxyl Radical Protein Footprinting and Protection Factor Analysis

NTD samples were purified into a buffer of 10 mM sodium phosphate, 0.05mM EDTA and pH 7.4. The protein concentration was adjusted to 5 μM. Samples were exposed to X-ray white beam for 0-50 ms at the 3.2.1 beamline of Advanced Light Source (Berkeley, CA) at ambient temperature, and immediately quenched with 10 mM methionine amide to prevent secondary oxidation. Subsequently, all samples were reduced with 10 mM DTT at 56 °C for 45 minutes and alkylated with 25 mM iodoacetamide at room temperature in the dark for 45 minutes. Two sets of protease digestion were performed at 37 °C using an enzyme-to-protein ratio of 1:10 (w/w). One was with Lys-C (Promega, Inc.) overnight, followed by Asp-N (Promega, Inc.) for 5 hours. The other was with pepsin (Promega, Inc.) for 2 hours. Digestion was terminated by heating samples at 95 °C for 2 min. Identification and quantification of oxidative sites were performed by liquid chromatography-mass spectrometry (LC-MS) analysis using an Orbitrap Elite mass spectrometer (Thermo Scientific, CA) interfaced with a Waters nanoAcquity UPLC system (Waters, MA). The proteolytic peptides (150 ng) were loaded on a trap column (180 μm × 20 mm packed with C18 Symmetry, 5 μm, 100 Å; Waters, MA) to desalt and concentrate peptides. Peptide mixture was eluted on a reverse phase column (75 μm × 250 mm column packed with C18 BEH130, 1.7 μm, 130 Å; Waters, MA) using a gradient of 2 to 35% mobile phase B (0.1% formic acid and acetonitrile) vs. mobile phase A (100% water/0.1 % formic acid) for 60 minutes at 40 °C at a flow rate of 300 nL/min. Eluted peptides were introduced into the nano-electrospray source at a capillary voltage of 2.5 kV.

The extent of oxidation by MS was quantified as a function of X-ray exposure time. The curves were fit to an exponential decay function and the measured rate constants (*k*_fp_) were divided by a measure of their intrinsic reactivity with hydroxyl radicals (*K*_R_), thereby providing a residue-level protection factor (PF=*k*_R_/*k*_fp_) (Huang et al., 2015; Kaur et al., 2015). The PF calculation is available from the website http://www.theyanglab.org/protection.html. The log of the PF values provides an accurate surface topology map, where high logPF values reflect more protection for solvent and lower logPF values reflect greater solvent exposure.

### Conformational Library

A pool of 35,240 NTD structure candidates was generated from molecular dynamics simulations with an accumulative total time of 35-μs in a 1-ns recording frequency. To enhance conformational sampling, two advanced algorithms were used: Gaussian accelerated molecular dynamics (GaMD) (Miao et al., 2015) and replica exchange solute tempering (REST2) (Wang et al., 2011). First, a set of 25 GaMD trajectories (using the software AMBER16, each starting with a random configuration and lasting 1 μs) resulted in a total of 25 μs. Second, a set of 64 replicas in REST2 simulations, ranging from 300 K to 600 K were performed at the Argonne Leadership Computing Facility using the software NAMD as previously described (Jo and Jiang, 2015; Phillips et al., 2005). Each replica lasted 160 ns, which resulted in 10 μs. In both GaMD and REST2 simulations, the molecular force field of Amber ff99SB with a TIP3P water model was used (Hornak et al., 2006). Given that the force field for IDPs is still under active development (Best et al., 2014; Huang and MacKerell, 2018; Huang et al., 2017; Nerenberg and Head-Gordon, 2018; Piana et al., 2015; Song et al., 2017), the Amber ff99SB was used here for the purpose of pose generation only, with the help from the inclusion of all trajectories from different temperatures and nonequilibrium simulations. Hence, the accuracy of the force field used is beyond the scope here, and these conformations were combined and selected by comparison with multi-technique experimental measurements. To group similar structures together and reduce the computational cost, we performed an RMSD-based clustering analysis as previously described (Huang et al., 2016; Svergun, 1992). Using a Ca-RMSD cutoff of 5 Å, we obtained a set of N=8,491 clusters of NTD structures from the total of 35,240 simulation snapshots.

### Ensemble Structure Construction

Based on the library of NTD conformations generated above, data from SAXS and FP-PF measurements were combined for ensemble refinement. We selected the ensemble-structures that best-fit experimental SAXS and FP-PF data simultaneously, by a weighted average over the resulting *N* clusters of structures with a probability distribution {P_*i*_} = *P*_1_,···,*P*_*N*_ of each cluster (*i*), based on two unit-less functions χ^2^ and φ^2^. The χ^2^ is defined to measure the difference between the theoretical and experimental SAXS profile by (Yang et al., 2010),

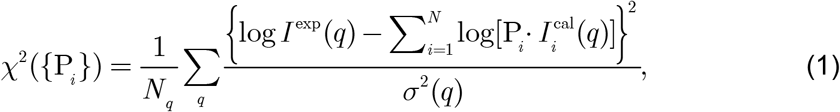

where *I*^cal^ is calculated using f*ast-SAXS-pro* (Ravikumar et al., 2013) and *N*_*q*_ is the number of scattering *q* points recorded in experimental *I*^exp^ (with its measurement error of σ(*q*)). Similarly, the φ^2^ is the goodness of fit between experimental and theoretical FP-FP data by (Hsieh et al., 2017; Huang et al., 2018; Huang et al., 2016),

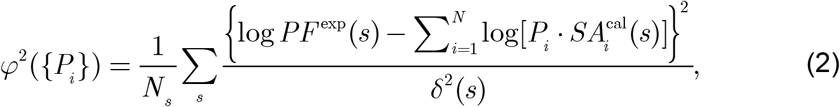

where SA^cal^ is the predicted side-chain SA value of each site, using the linear regression between experimental log*PF*^exp^ values (with an error *δ*(*s*) at each site (*s*)) and SA values of *N*_*s*_ sites for each structural cluster (here, *N*_s_=16 as shown in Table S3).

To avoid potential overfitting, we performed a maximum-likelihood principle analysis by adopting a “relative entropy” contribution (Boomsma et al., 2014; Rozycki et al., 2011),

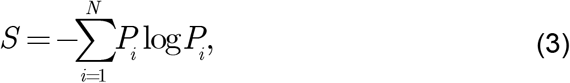

assuming a uniform prior, so the entire optimization of the probability distribution was achieved by minimizing an effective free energy,

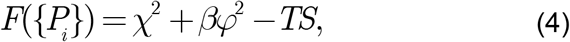

where *β* is a scaling factor for balancing two unit-less functions χ^2^ and ϕ^2^ and *T* is an effective temperature-like scaling factor. We chose *β* = 0.1 and *T* = 0.05 by minimizing *F*(*β,T*) (see Fig. S5). To reduce the computational cost, we decreased the number of clusters to Nβ519 by satisfying a cutoff criteria of χ^2^ < 5 and a negative linear correlation between logPF and SA for each cluster. Finally, an optimal probability distribution {*P*_*i*_} was determined by numerically minimizing Eq. (4) with the constraints of 0 <*P*_*i*_ < 1 and Σ_*i*_*P*_*i*_ = 1 using a genetic algorithm (Conn et al., 1997).

### Ensemble Contact Map

The residue-residue interaction (or contact) was calculated using the *contact of structural units* (CSU) approach (Sobolev et al., 1999). Each residue was the center of a window segment of 5 neighboring amino acids. Each contact was weighted over the ensemble-structures by the probability of top 100 structures that account for a total of 80% of the entire population.

### Surface Plasmon Resonance

The binding of NTD with TBP was measured in PBSP+ (GE healthcare) buffer, pH7.4 by surface plasmon resonance (SPR) using a Biacore T100 system (GE Healthcare) at 4 °C. S series sensor chip CM4 (GE Healthcare) was preconditioned at 100uL/min with successive 20 μL pulses of 50 mM NaOH, 100 mM HCl, 0.1 mM SDS and 0.085% (v/v) H_3_PO_4_. NTD was immobilized at low density and TBP concentration series ranging from 1.5 to 25 μM were injected over the NTD surface. Data were plotted by Origin 2017 and the affinity constant was calculated by Biaevaluation3 software (GE Healthcare).

### Fluorine-19 NMR Spectroscopy

The NTD^S118C^-BTFA protein was prepared by site-directed mutation of S118C and BTFA (3-Bromo-1,1,1-trifluoroacetone, Alfa Aesar Cat# A14948) was added during the solubilization of inclusion bodies with a 1000-fold dilution and further incubated overnight at 4 °C. The final NTD-BTFA protein was exchanged into a buffer of 10mM sodium phosphate, pH 7.2, 0.5 mM EDTA, and 0.1 mM PMSF. The final protein concentration used for ^19^F-NMR is 2.0 mg/ml, 1.6 mg./ml, and 2.1 mg/ml for the wild-type, L31A, the triple mutant K32A/I33A/P34A of the NTD^S118C^-BTFA, respectively. A volume of 250 μl was used for each protein sample containing 10% D_2_O as an initial point for D_2_O titration from 10% to 50%.

One-dimensional ^19^F-NMR experiments were performed on a 500 MHz Bruker Ascend III HD spectrometer equipped with a nitrogen-cooled ^19^F tuned BBO probe. Spectra were acquired with 512 scans, 131K data points in the direct dimension, a pulse length of 15.0 μsec, a spectral width 468750 Hz (^19^F), a digital resolution of 3.5 Hz/point, and a relaxation delay of 1.0 s at the temperature of 281 K. The data was processed with Topspin 3.2 software and free induction decay (FID) signals were apodized with an exponential window function as to line broadening of 0.30 Hz.

### >Quantification and Statistical Analysis

Duplicates were performed for protein footprinting and standard deviations were indicated. SAXS data uncertainties were derived from a total of five scattering images. The ensemble fitting calculations were repeated nine times; reported probability values are the average and standard deviation for the replicated calculations.

### Data and Software Availability

The SAXS data and three representative structures for the NTD ensemble derived from inhouse multi-technique modeling have been deposited in the Small Angle Scattering Biological Databank (SASBDB access code SASDEE2; https://www.sasbdb.org/data/SASDEE2). A set of 10 integrative ensemble-structures of the ERα-NRD have been deposited to the multi-scale I/H structure PDB-dev database (PDB-dev access code PDBDEV_00000027; https://pdb-dev.wwpdb.org/static/cif/PDBDEV_00000027.cif). Other data are available from the corresponding author upon reasonable request.

---

**Figure S1.**
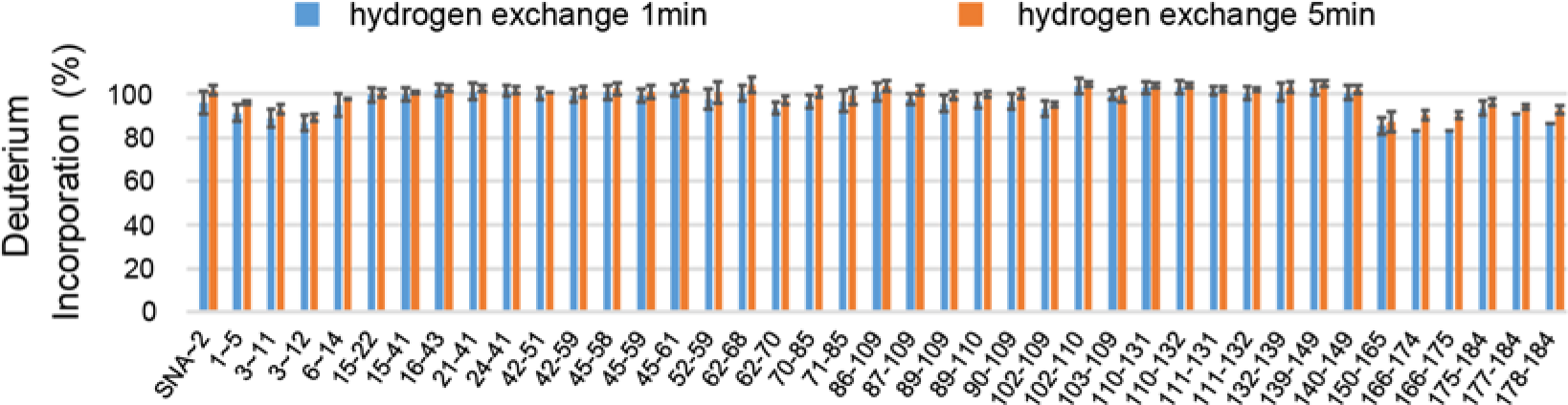
Deuterium incorporation of peptic fragments in HX-MS within 1 min (blue) and 5 min (red) incubation in D_2_O. Error bars indicate standard deviation (three independent experiments).

**Figure S2.**
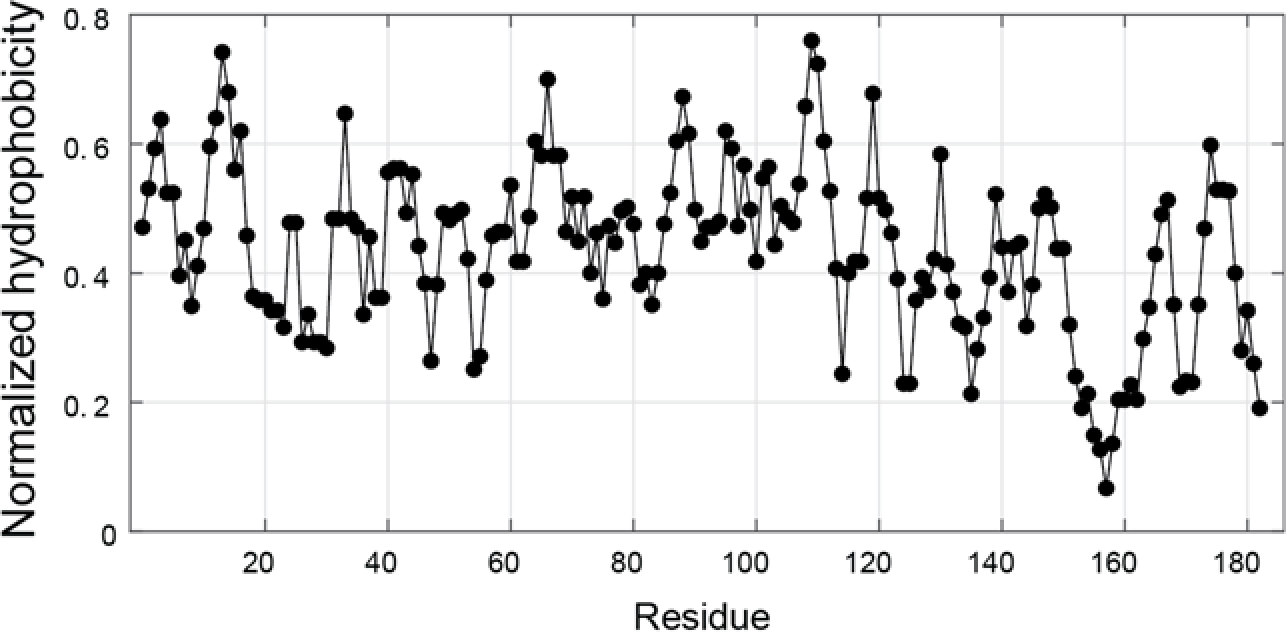
The hydrophobicity of amino acids in the NTD sequence. Each was calculated by the Kyte-Doolittle approximation using the ExPASy server (with a window size of 5 amino acids). The hydrophobicity of individual residues was normalized to a scale of 0 to 1. The mean hydrophobicity <*H*> is defined as the sum of the normalized hydrophobicities of all residues divided by the number of residues in the polypeptide, yielding a <*H*> value at 0.446.

**Figure S3.**
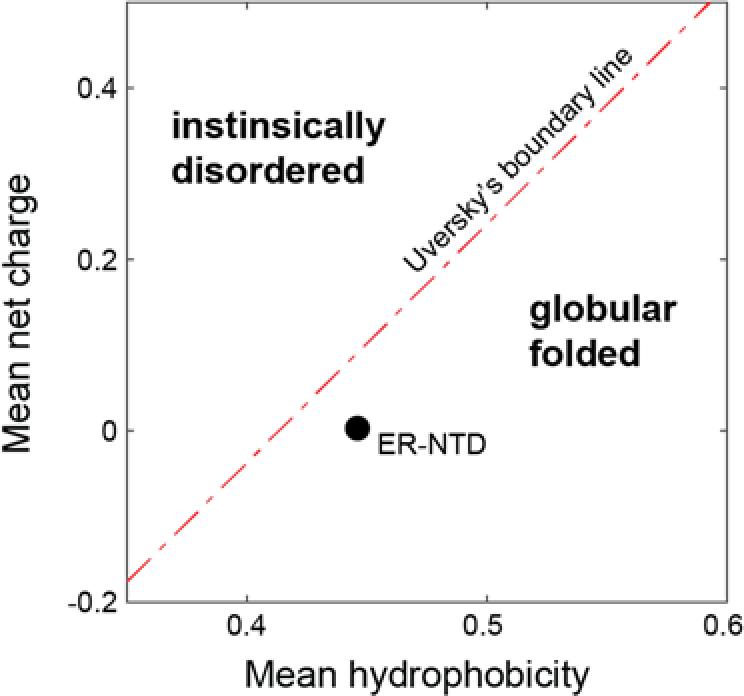
The two-dimensional space of mean hydrophobicity <*H*> and mean net charge <*q*> of globular and intrinsically disordered proteins. The ERα-NTD is close to the Uversky’s boundary line of <*H*> = (<*q*> + 1.151)/2.785, and enters the globular-fold regime.

**Figure S4.**
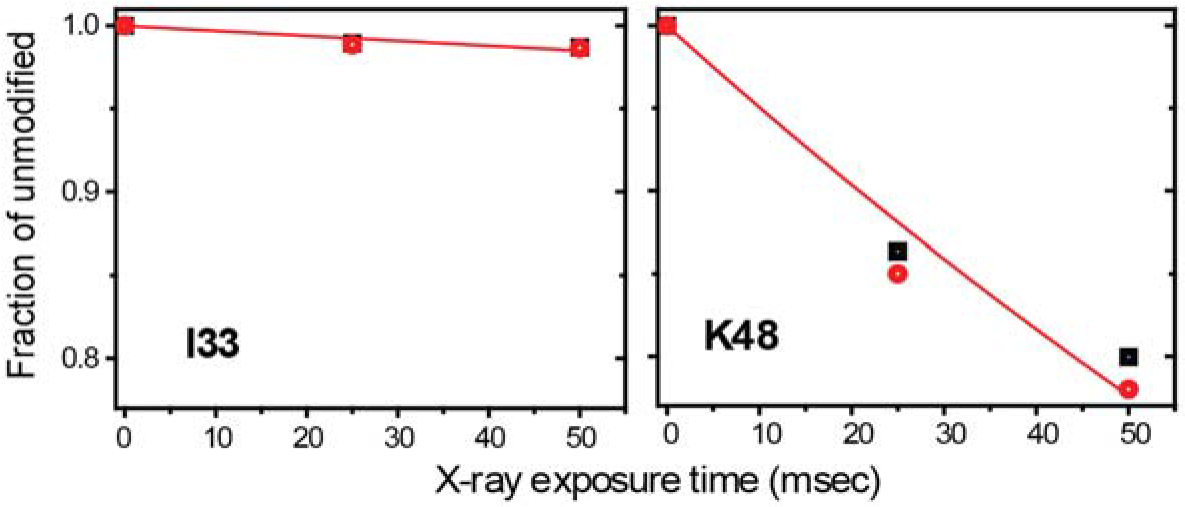
Hydroxyl radical footprinting dose-response curves as a function of X-ray exposure time. Red lines are the least-squares fit to an exponential decay function (i.e., a normalized fraction of unmodified residues), each yielding a rate of footprinting (*k*_fp_). The division by an intrinsic reactivity with hydroxyl radicals (*k*_R_) provides a protection factor measure (PF = *k*_R_/*k*_fp_).

**Figure S5.**
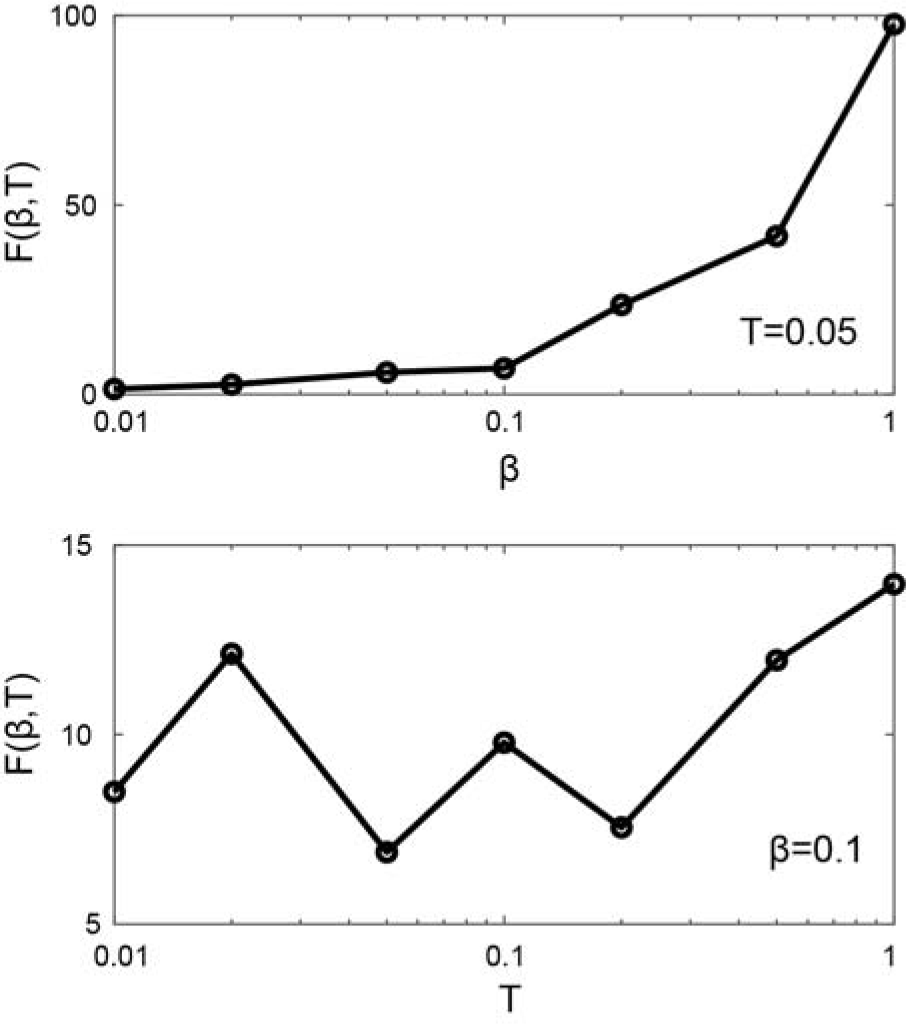
The effective free energy F(β,T) as function of the parameters β and T. The values of β=0.1 and T=0.05 were chosen to give the local minimum of F(β,T) (see Eq. 4).

**Figure S6.**
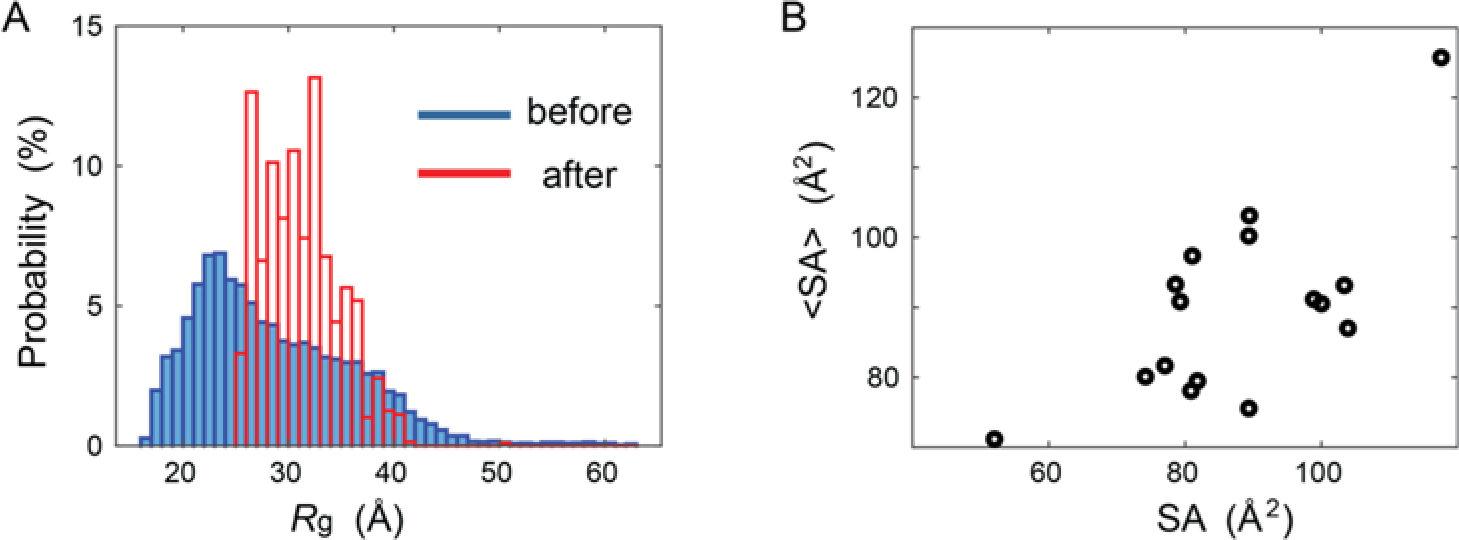
Structure selection by the ensemble-fitting method against experimental scattering and footprinting data. (A) The histogram plot as a function of *R*_g_ before and after the ensemble fitting was applied. (B) The ensemble-averages of solvent-accessible surface area for the set of 16 footprinting-probed residues before (<SA>) and after (SA) the ensemble-fitting.

**Figure S7.**
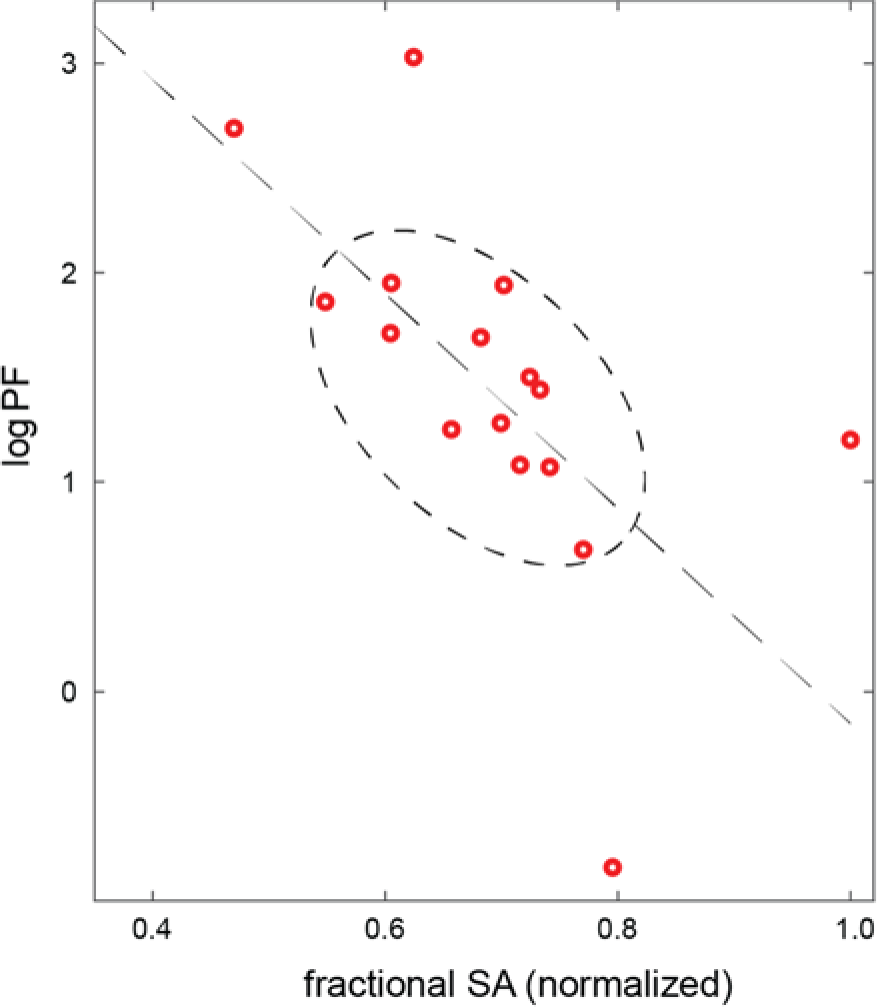
Measured logPF values against the fractional SA values for the set of 16 probed residues. The factional SA value was the SA weighted by each residue’s maximum possible accessible area.

**Figure S8.**
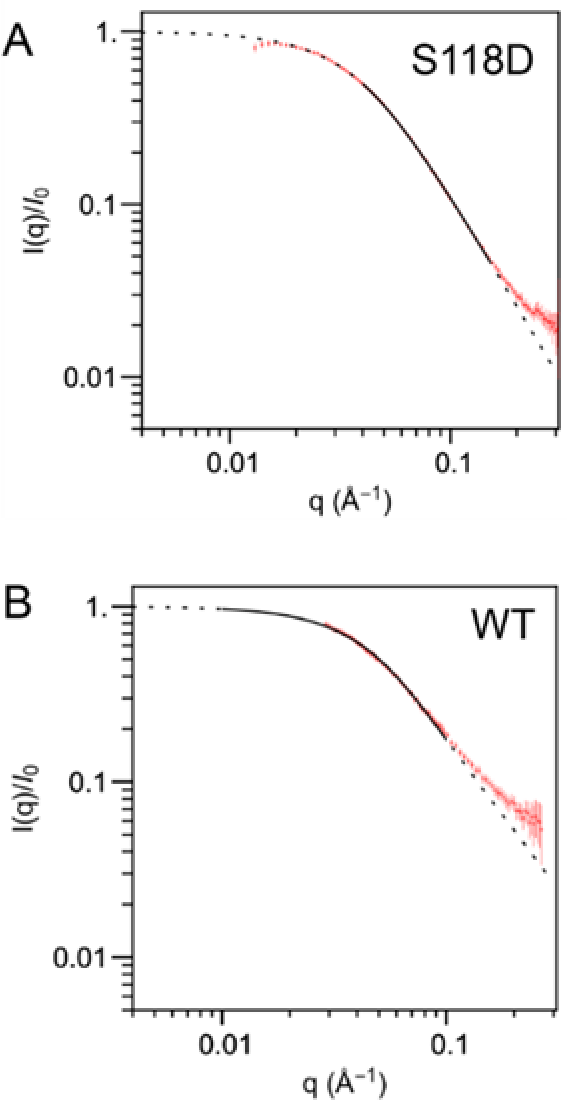
SAXS data for mutant S118D (A), compared to the wild-type (B). Generated using the webserver at http://sosnick.uchicago.edu/SAXSonIDPs.

**Figure S9.**
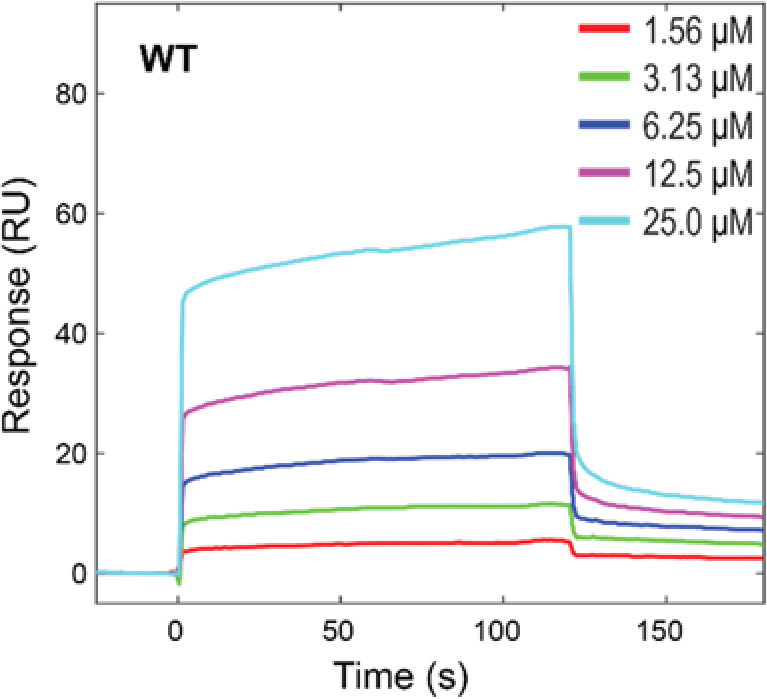
Binding of NTD with TBP measured by surface plasmon resonance (SPR) spectrometry. An apparent binding affinity was estimated at K_*d*_ = 53 ± 12 μM.

**Figure S10.**
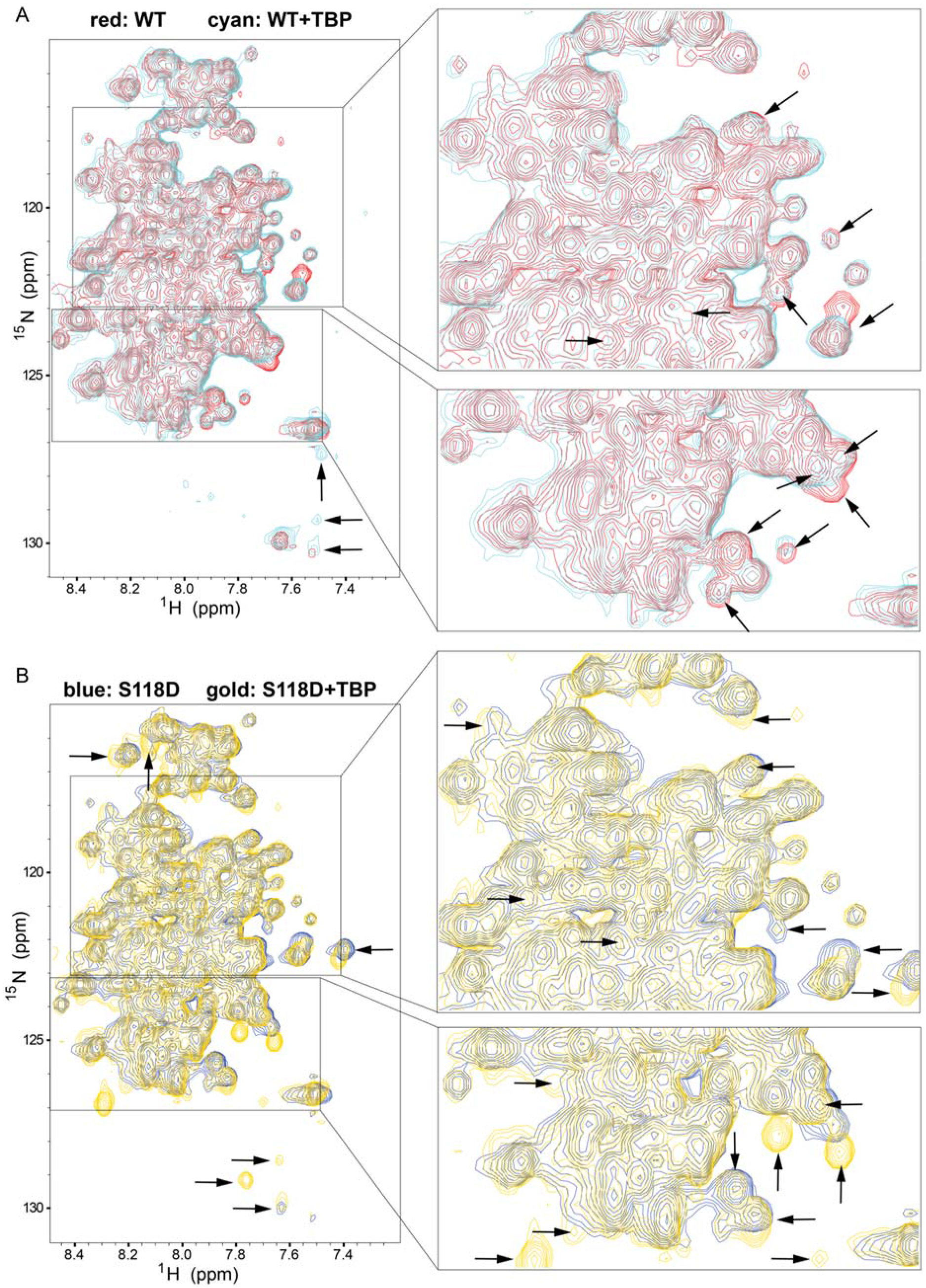
Mutation-induced conformational changes and alteration of coactivator TBP binding. (A) Overlap of ^1^H-^15^N TROSY spectra of the WT in the absence (red) and presence (cyan) of TBP. Change/shift of peaks is marked by arrows. (B) Overlap of ^1^H-^15^N TROSY spectra of S118D in the absence (blue) and presence (gold) of TBP.

**Figure S11.**
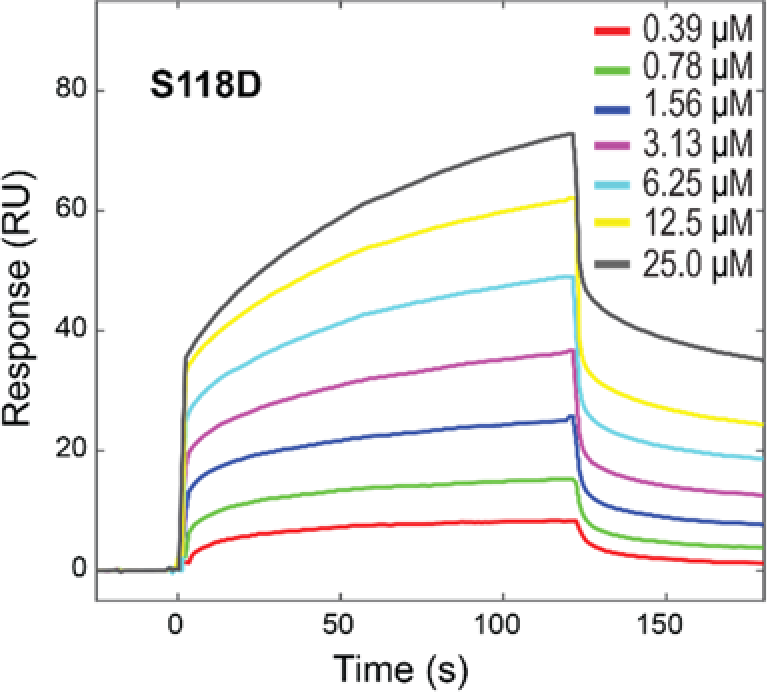
Measured TBP-binding for the S118D mutant using SPR spectrometry. An apparent binding affinity was estimated at K_*d*_ = 3.9 ± 0.4 μM.

**Table S1.** SAXS data details and parameters.

|  |  |
| --- | --- |
| Experiment date | June 12, 2017 |
| SASBDB ID<br>(https://www.sasbdb.org) | SASDEE2<br>(https://www.sasbdb.org/data/SASDEE2) |
| Beamline/Instrument | NSLS-II/16-ID-LiX, Pitaus3 S 1M/300k |
| Wavelength | 0.9 Å |
| Sample detector distance | 4.270 m / 0.856 m |
| Exposure time | 1 second |
| Cell temperature | 10 °C |
| protein concentration | 2.5 mg/ml |
| Number of residues | 187 |
| Molecular weight | 20.18 kDa |
| Protein buffer | 20mM Hepes, pH7.4, 300mM NaCl, 5%<br>glycerol, 1mM DTT, 0.5 mM PMSF, 1 mM<br>EDTA |

**Table S2.** Radius of gyration (*R*_g_) values derived from SAXS data using several fitting methods.

| WT/S118D | Guinier | Debye | extended Guinier | Sosnick |
| --- | --- | --- | --- | --- |
| $R_g$ (Å) | 30.0±1.6/35.8±0.5 | 32.9± 0.6/41.0±0.3 | 29.2±1.3/36.7±0.6 | 31.0±0.2/38.7±0.2 |
| $(qR_g)_{\max}$ | 1.3 | 3.0 | 2.0 | 3.0/5.2 |
| reference | (Guinier, 1939) | ( Debye,<br>1947;Calmettes et al.,<br>1994) | (Zheng and Best, 2018) | (Riback et al., 2017) |

**Table S3.**
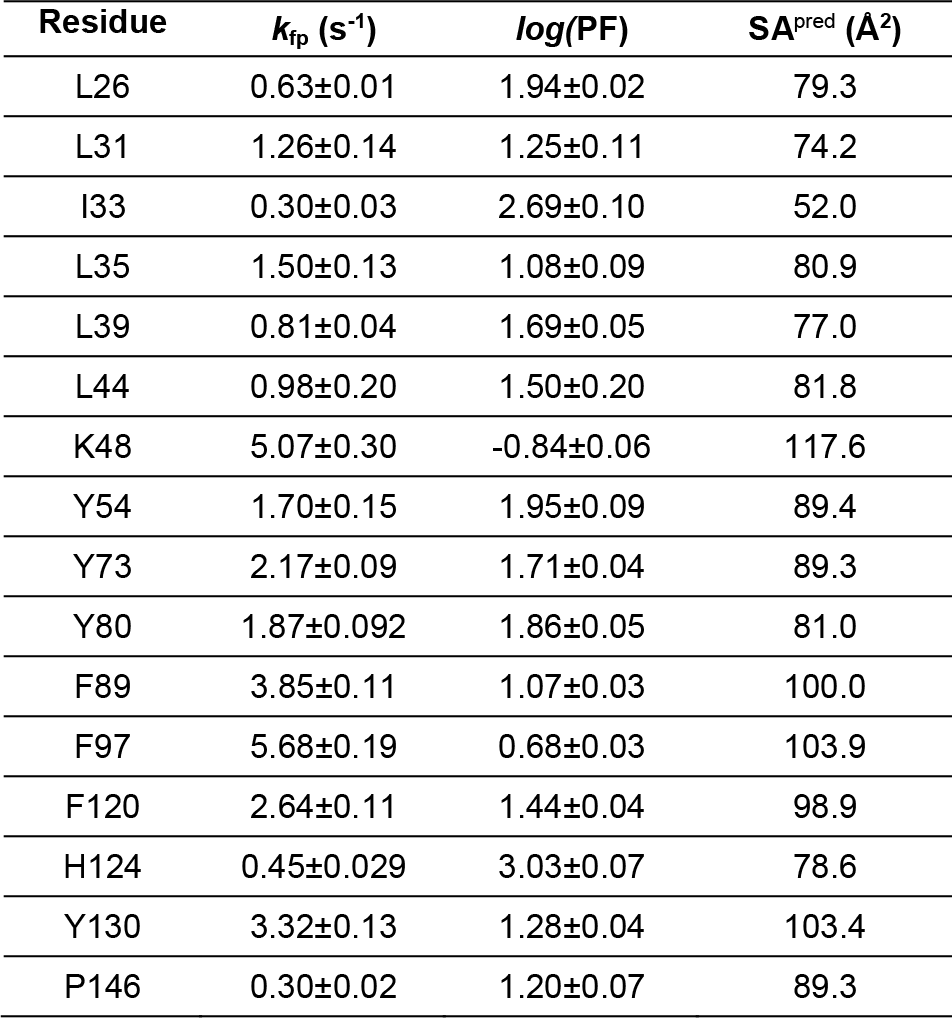
Hydroxyl radical footprinting protection factors (FP-PFs) for the NTD residues. Listed are the probed residues and their corresponding values for footprinting rates (k_fp_), logPF, and SA^pred^ predicted from the ensemble-structures.

| Residue | $k_{fp}$ ($s^{-1}$) | log(PF) | $SA^{pred}$ ($\text{\AA}^2$) |
| --- | --- | --- | --- |
| L26 | 0.63±0.01 | 1.94±0.02 | 79.3 |
| L31 | 1.26±0.14 | 1.25±0.11 | 74.2 |
| I33 | 0.30±0.03 | 2.69±0.10 | 52.0 |
| L35 | 1.50±0.13 | 1.08±0.09 | 80.9 |
| L39 | 0.81±0.04 | 1.69±0.05 | 77.0 |
| L44 | 0.98±0.20 | 1.50±0.20 | 81.8 |
| K48 | 5.07±0.30 | -0.84±0.06 | 117.6 |
| Y54 | 1.70±0.15 | 1.95±0.09 | 89.4 |
| Y73 | 2.17±0.09 | 1.71±0.04 | 89.3 |
| Y80 | 1.87±0.092 | 1.86±0.05 | 81.0 |
| F89 | 3.85±0.11 | 1.07±0.03 | 100.0 |
| F97 | 5.68±0.19 | 0.68±0.03 | 103.9 |
| F120 | 2.64±0.11 | 1.44±0.04 | 98.9 |
| H124 | 0.45±0.029 | 3.03±0.07 | 78.6 |
| Y130 | 3.32±0.13 | 1.28±0.04 | 103.4 |
| P146 | 0.30±0.02 | 1.20±0.07 | 89.3 |

